# Liver fibrosis uncouples tumor control from survival after multikinase inhibitor immunotherapy in hepatocellular carcinoma

**DOI:** 10.1101/2023.10.20.563378

**Authors:** Satoru Morita, Tomofumi Ando, Hiroto Kikuchi, Atsuyo Morita, Tatsuya Kobayashi, Grace Birch, Ryota Tanaka, Aya Matsui, Zhiping Ruan, Peigen Huang, Yeonju Cho, Alexei Hernandez, Jae W Lee, Erin M. Coyne, Sarah M. Shin, Robin K. Kelley, Mark Yarchoan, Stefan Halvorsen, Slim Sassi, Mari Mino-Kenudson, Shadmehr Demehri, Rizwan Romee, Won Jin Ho, Dan G. Duda

## Abstract

Combining multikinase inhibitors with immune checkpoint blockade improves tumor control in hepatocellular carcinoma (HCC), but survival benefits remain inconsistent. To define this discordance, we integrated COSMIC-312 with orthotopic and autochthonous murine HCC models with or without liver fibrosis. In patients treated with cabozantinib plus atezolizumab, baseline liver function stratified overall survival but not progression-free survival, indicating uncoupling of tumor control from survival. In contrast, sorafenib outcomes tracked with both endpoints, supporting a treatment-specific effect rather than a purely prognostic effect of liver status. In murine HCC models, cabozantinib plus PD-1 blockade induced comparable tumor regression regardless of liver condition, but improved survival only in mice with preserved liver function, whereas fibrotic hosts developed hepatotoxicity. Immune profiling revealed compartment-specific remodeling, with enhanced cytotoxic T-cell programs in tumors but expansion of NK-lineage innate lymphocytes with ILC1-like features in fibrotic liver. Depletion of NK1.1-positive cells reduced liver injury and restored survival without compromising antitumor efficacy, whereas CD4-positive or CD8-positive T-cell depletion did not protect from hepatotoxicity. Transcriptomic, single-cell, adoptive-transfer, and human in vitro studies supported a model in which the fibrotic liver niche promotes NK-to-ILC1-like reprogramming, hepatocyte stress signaling, and TNF/TRAIL-associated epithelial injury. Consistently, cabozantinib and nivolumab showed liver-predominant remodeling of CD56-positive innate lymphocyte-enriched populations in human HCC samples, and ex vivo TGF-beta induced ILC1-like phenotypic changes in human NK cells. These findings identify liver fibrosis as a host determinant that can limit the survival benefit of multikinase inhibitor immunotherapy by promoting innate immune-mediated hepatotoxicity despite preserved tumor control in HCC.

**One Sentence Summary:** Liver fibrosis limits the survival benefit of multikinase inhibitor immunotherapy in hepatocellular carcinoma by promoting innate immune–mediated hepatotoxicity despite preserved tumor control.

## INTRODUCTION

Hepatocellular carcinoma (HCC) is a leading cause of cancer-related mortality worldwide and most commonly arises in the setting of chronic liver disease and cirrhosis. Because curative therapies are feasible in only a minority of patients, systemic treatment remains the mainstay for advanced disease. Immune checkpoint blockade (ICB) has transformed the treatment of multiple malignancies, including advanced HCC, but responses to monotherapy remain limited, prompting the development of combination strategies to improve efficacy. In HCC, however, therapeutic benefit may depend not only on tumor control but also on preservation of host liver function.

Immunotherapy-based combinations have reshaped first-line therapy for advanced HCC. In particular, anti-PD-L1 plus anti-VEGF antibody therapy improves survival and has established the principle that modulating both tumor immunity and the vascular microenvironment can enhance clinical benefit (*1*). This success motivated the evaluation of multikinase inhibitors (MKIs) with ICB, given their capacity to target angiogenic and oncogenic pathways while also altering immune responses within tumors and peripheral tissues. In the randomized phase III COSMIC-312 trial, cabozantinib plus atezolizumab significantly prolonged progression-free survival (PFS) compared with sorafenib but failed to improve overall survival (OS) (*2*). This discordance between tumor control and survival remains unexplained and raises the possibility that, in HCC, host organ context can limit the benefit of immunotherapy-based combinations independently of tumor response.

This question is especially relevant in HCC because most patients have underlying liver fibrosis or cirrhosis (*3*). As a result, survival is determined not only by tumor progression but also by hepatic reserve and susceptibility to treatment-related injury. A key unresolved issue is whether combination therapies that enhance antitumor immunity may also narrow the therapeutic index by exacerbating injury to the non-malignant fibrotic liver. The dissociation between PFS and OS observed in COSMIC-312 suggests that pre-existing liver damage may selectively compromise treatment tolerability or post-progression fitness without necessarily reducing on-tumor efficacy. More broadly, this framework could help explain why several MKI+ICB strategies in HCC have yielded tumor responses without commensurate survival benefit.

The liver contains abundant populations of group 1 innate lymphocytes, including blood-derived conventional natural killer (cNK) cells and tissue-resident type 1 innate lymphoid cells (ILC1), which contribute to tissue surveillance and homeostasis (*4–6*). Under conditions of chronic injury and fibrosis, these populations can acquire pathogenic functions and produce inflammatory cytokines and death ligands such as tumor necrosis factor (TNF) and TNF-related apoptosis-inducing ligand (TRAIL), both of which can promote hepatocyte damage (*7–9*). Moreover, NK-cell plasticity in TGF-β-rich environments can drive acquisition of ILC1-like features, suggesting a mechanism by which the fibrotic liver niche might reshape innate immunity during therapy (*10–12*). However, whether innate lymphocyte reprogramming contributes to treatment-associated hepatotoxicity and limits survival benefit from MKI-ICB combinations in HCC remains unknown.

Here, we investigated the basis of the discordance between tumor control and survival after MKI-ICB therapy in HCC. We analyzed clinical outcomes from the randomized phase III COSMIC-312 trial according to baseline liver function and complemented these studies with orthotopic and autochthonous murine HCC models with or without pre-existing liver fibrosis and analyses of tissues from advanced HCC patients treated with cabozantinib and nivolumab (*13*). Using integrated immune profiling, transcriptomic analysis, adoptive-transfer experiments, and human translational studies, we examined how liver fibrosis shapes intrahepatic immune responses, hepatotoxicity, and therapeutic outcomes. We hypothesized that liver fibrosis narrows the therapeutic index of MKI-ICB therapy by promoting a hepatotoxic innate immune state that limits survival benefit despite preserved tumor control.

## RESULTS

### Baseline liver function stratifies survival but not tumor control after cabozantinib/ICB

To test whether host liver status contributes to the discordance between tumor control and survival after immunotherapy-based treatment in HCC, we analyzed outcomes from the randomized phase III COSMIC-312 trial according to baseline liver function (**Fig. 1A-D, Fig. S1A-H and Tables S1-S3**). In patients treated with cabozantinib plus atezolizumab, baseline liver function stratified overall survival (OS) but not progression-free survival (PFS): patients with preserved liver function (APRI<0.7) had significantly longer OS than those with impaired liver status, whereas PFS was not significantly different between groups (**Fig. 1A, B**). In contrast, among sorafenib-treated patients, liver function tracked with both OS and PFS, indicating a more conventional coupling of tumor progression and survival (**Fig. S1A, B**).

**Fig. 1.**
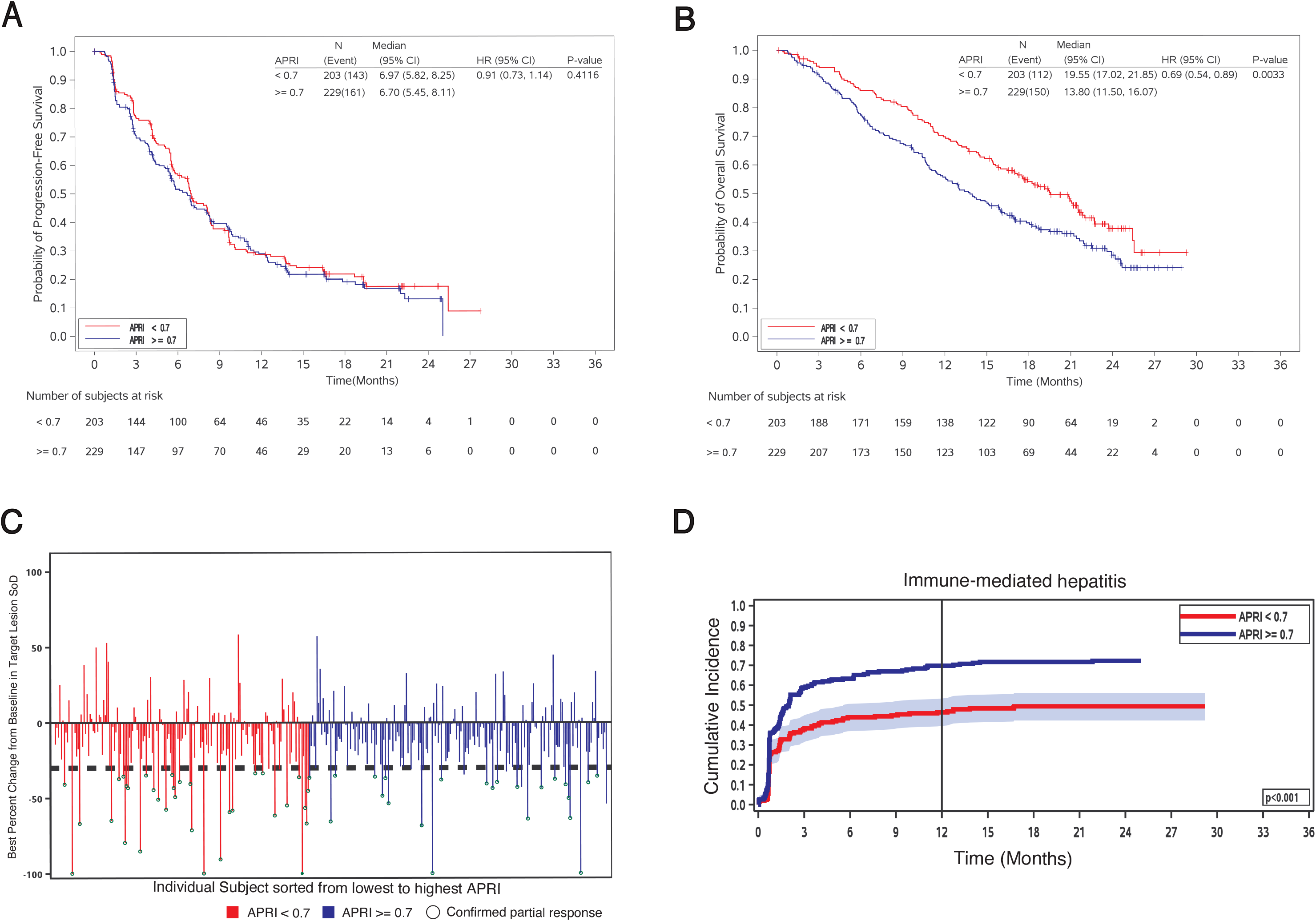
Baseline liver dysfunction uncouples survival from tumor control and increases reported liver-related toxicity after cabozantinib/atezolizumab therapy in advanced HCC. (A, B) Kaplan-Meier curves for progression-free survival (PFS) (A) and overall survival (OS) (B) in patients with advanced hepatocellular carcinoma treated with cabozantinib plus atezolizumab in the COSMIC-312 trial, stratified by baseline aspartate aminotransferase-to-platelet ratio index (APRI <0.7 versus APRI >=0.7). Hazard ratios (HRs), 95% confidence intervals (CIs), median survival, P values, and numbers at risk are shown. (C) Waterfall plot showing best percent change from baseline in the sum of diameters of target lesions in cabozantinib/atezolizumab-treated patients, ordered from lowest to highest baseline APRI. Bars indicate APRI strata, and open circles indicate confirmed partial responses. (D) Cumulative incidence of reported immune-mediated hepatitis/liver-related toxicity in cabozantinib/atezolizumab-treated patients stratified by baseline APRI. P values are shown in the panels.

As an exploratory sensitivity analysis, we modeled APRI as a continuous variable using restricted cubic splines (**Fig. S1C-F**). These analyses were directionally consistent with the primary categorical APRI findings, showing no clear relationship between APRI and PFS in the cabozantinib/atezolizumab arm, while the OS relationship was less suitable for visual dose-response interpretation because of broad confidence intervals at higher APRI values. We therefore used the spline analysis only as supportive evidence and based our primary clinical inference on prespecified APRI strata, APRI-ALBI validation, tumor-reduction analysis, competing-risk toxicity analysis, and multivariable modeling. Maximum tumor reduction occurred across the full range of APRI values in cabozantinib/ICB-treated patients, indicating preserved radiographic tumor control despite impaired liver function (**Fig. 1C**). Competing-risk analysis showed a higher incidence of liver-related toxicity in patients with worse baseline liver status, consistent with increased susceptibility of fibrotic livers to treatment-associated injury **(Fig. 1D**).

These findings were reproduced using a combined APRI-ALBI metric and remained significant after multivariable adjustment for major clinical covariates (**Fig. S1G, H and Tables S1, S2**) (14, 15). Patients with impaired liver function were also less likely to receive subsequent systemic therapy and had shorter post-protocol treatment trajectories, consistent with early clinical deterioration despite initial tumor control (**Table S3**). Together, these data indicate that baseline liver dysfunction selectively limits survival benefit from cabozantinib/ICB without proportionally reducing tumor control, revealing treatment-specific uncoupling of response from outcome (**Fig. 1A-D and Fig. S1A-H**).

### Cabozantinib-containing regimens are associated with liver injury signals in clinical datasets

Because impaired liver status preferentially affected survival in the cabozantinib/ICB arm, we next asked whether cabozantinib exposure is associated with liver injury signals in independent clinical datasets (**Fig. S2A-C and Tables S4-S6**). Pharmacovigilance analyses showed enrichment of hepatobiliary adverse-event reporting among patients receiving cabozantinib-containing regimens (**Fig. S2A, B**). In an independent real-world cohort, cabozantinib-treated patients exhibited more frequent liver enzyme abnormalities than sorafenib-treated patients after propensity matching (**Fig. S2C and Tables S4, S5**). Across published trials, transaminase elevations were also more common in cabozantinib-containing arms (**Table S6**).

Because these datasets cannot definitively distinguish direct drug-induced liver injury from immune-mediated hepatitis, these analyses were interpreted conservatively as evidence of increased hepatotoxicity signals rather than proof of a single injury mechanism. Nonetheless, together with the COSMIC-312 stratification, these findings support the hypothesis that cabozantinib-based therapy imposes a clinically meaningful burden on the injured liver (**Fig. S2A-C, Tables S4-S6**).

### Pre-existing fibrosis uncouples tumor response from survival in murine HCC

To test causality and define mechanism, we modeled cabozantinib-based therapy in orthotopic murine HCC with either normal or fibrotic liver microenvironments (**Fig. 2A-D and Fig. S3A-H**). In mice bearing orthotopic RIL-175 tumors in the normal liver, cabozantinib plus anti-PD-1 induced marked tumor regression and significantly prolonged survival relative to control or monotherapy groups (**Fig. 2A, B and Fig. S3A**). Similar antitumor activity and survival benefit were observed in a metastatic and relatively ICB-resistant HCA-1 model, confirming efficacy when baseline liver condition was preserved (**Fig. S3B, C**).

**Fig. 2.**
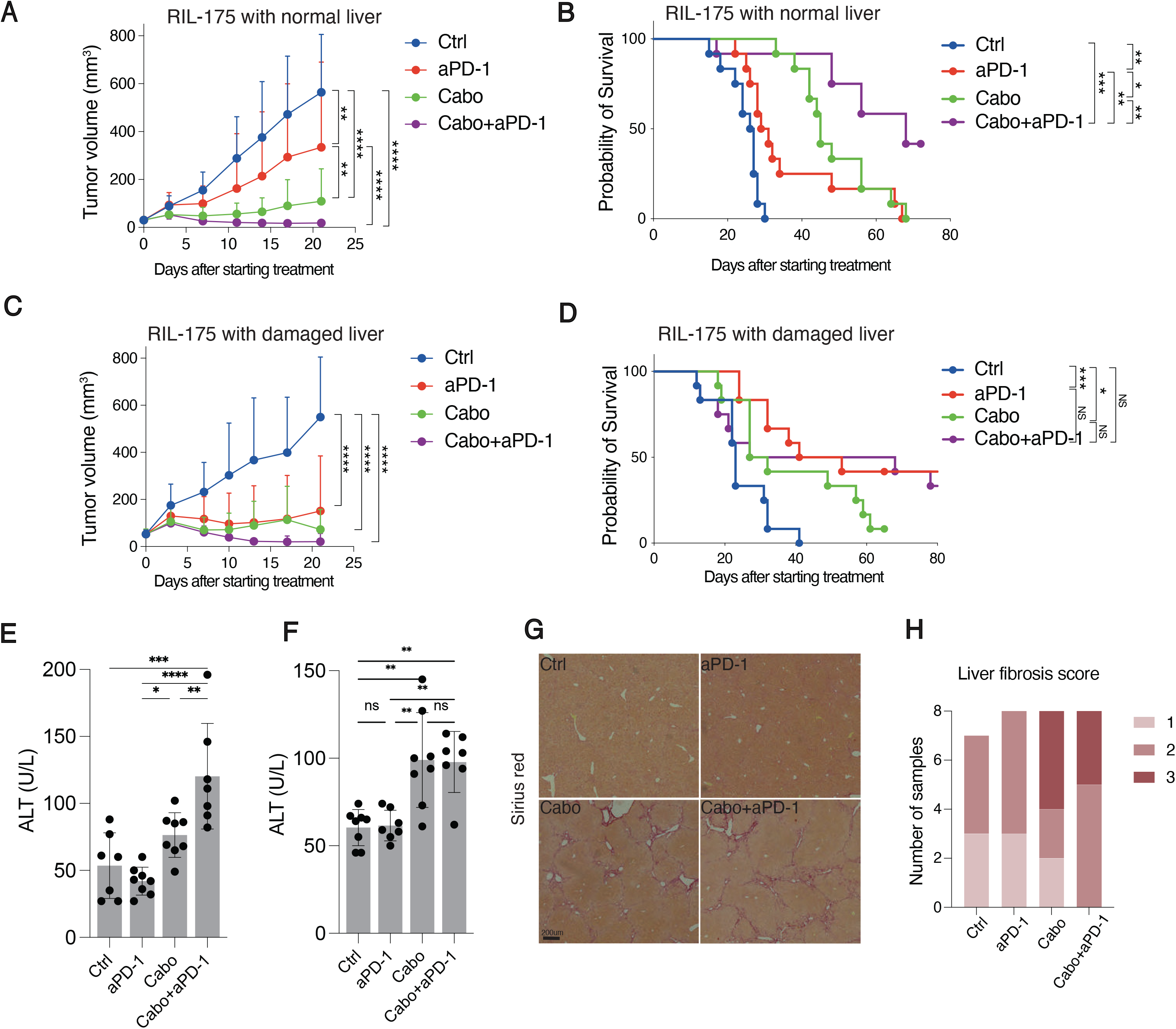
Cabozantinib/PD-1 blockade preserves antitumor activity but loses survival benefit and exacerbates liver injury in fibrotic murine HCC. (A, B) Tumor growth kinetics (A) and Kaplan-Meier survival curves (B) in mice bearing orthotopic RIL-175 HCC in normal liver treated with control, anti-PD-1 antibody, cabozantinib, or cabozantinib plus anti-PD-1. (C, D) Tumor growth kinetics (C) and Kaplan-Meier survival curves (D) in mice bearing orthotopic RIL-175 HCC in CCl_4_-induced fibrotic liver treated with the same regimens. (E, F) Plasma alanine aminotransferase (ALT) levels after treatment in RIL-175 tumor-bearing mice with fibrotic liver (E) and in an autochthonous Mst1^−/−^Mst2^f/–^ HCC model with liver fibrosis (F). (G, H) Representative Sirius red staining of surrounding non-tumor liver tissue (G) and liver fibrosis score distribution (H) after treatment in fibrotic hosts. Scale bar, 200 μm. Data are presented as mean +/- SD where applicable. *P < 0.05, **P < 0.01, ***P < 0.001, ****P < 0.0001; NS, not significant.

We next induced pre-existing fibrosis with CCl_4_ before tumor implantation (*16*) (**Fig. 2C, D and Fig. S3D-H**). In this setting, cabozantinib/ICB retained potent antitumor activity, with tumor growth inhibition and regression comparable to those seen in mice with normal liver (**Fig. 2C and Fig. S3D-F**). This preserved antitumor activity was also observed in the autochthonous HCC model with liver fibrosis, in which cabozantinib-containing therapy reduced relative tumor burden and Ki67-positive tumor-cell proliferation (**Fig. S3E, F**). However, the survival benefit was lost in fibrotic hosts despite preserved tumor control, and this dissociation was reproduced across orthotopic, metastatic, and autochthonous models(*16*) (**Fig. 2D and Fig. S3E-H**). These results mirror the clinical pattern observed in COSMIC-312 and demonstrate that host liver condition, rather than loss of antitumor efficacy, determines whether tumor response translates into survival benefit (**Fig. 2A-D, Fig. S3A-H**).

### Cabozantinib-based therapy selectively exacerbates hepatotoxicity in fibrotic liver

We next examined whether the loss of survival benefit in fibrotic hosts was accompanied by treatment-related liver injury (**Fig. 2E-H and Fig. S3I-K**). In mice with pre-existing fibrosis, both cabozantinib monotherapy and cabozantinib/ICB induced significant body weight loss, whereas mice with normal livers tolerated treatment without comparable systemic toxicity (**Fig. S3I, J**). A similar body-weight loss pattern was observed in the autochthonous HCC model with liver fibrosis, supporting the reproducibility of treatment-associated toxicity in an independent fibrosis-associated HCC model (**Fig. S3K**). Plasma ALT levels increased significantly after cabozantinib-containing therapy in fibrotic hosts but not in non-fibrotic hosts (**Fig. 2E, F**). Histologic analyses further demonstrated increased fibrosis and injury in surrounding non-tumor liver tissue after treatment (**Fig. 2G, H**).

Thus, pre-existing fibrosis did not reduce tumor response but instead narrowed the therapeutic index by increasing off-tumor hepatotoxicity (**Fig. 2C-H and Fig. S3I-K**). These findings provide a mechanistic framework for the clinical uncoupling of tumor control from survival after cabozantinib/ICB (**Fig. 1**, **Fig. 2**).

### Cabozantinib/ICB induces compartment-specific immune remodeling in tumor and fibrotic liver

To define immune mechanisms underlying this dissociation, we performed CyTOF profiling of orthotopic RIL-175 tumors and matched surrounding liver tissue after treatment (**Fig. 3A-G** and **Fig. S4A-H**). Unsupervised clustering revealed compartment-specific immune remodeling rather than a uniform systemic response (**Fig. 3A, B** and **Fig. S4A**). Within tumors, cabozantinib/ICB increased cytotoxic CD8-positive T-cell states and reduced suppressive programs, consistent with preserved or enhanced antitumor immunity (**Fig. 3C** and **Fig. S4B, C**).

**Fig. 3.**
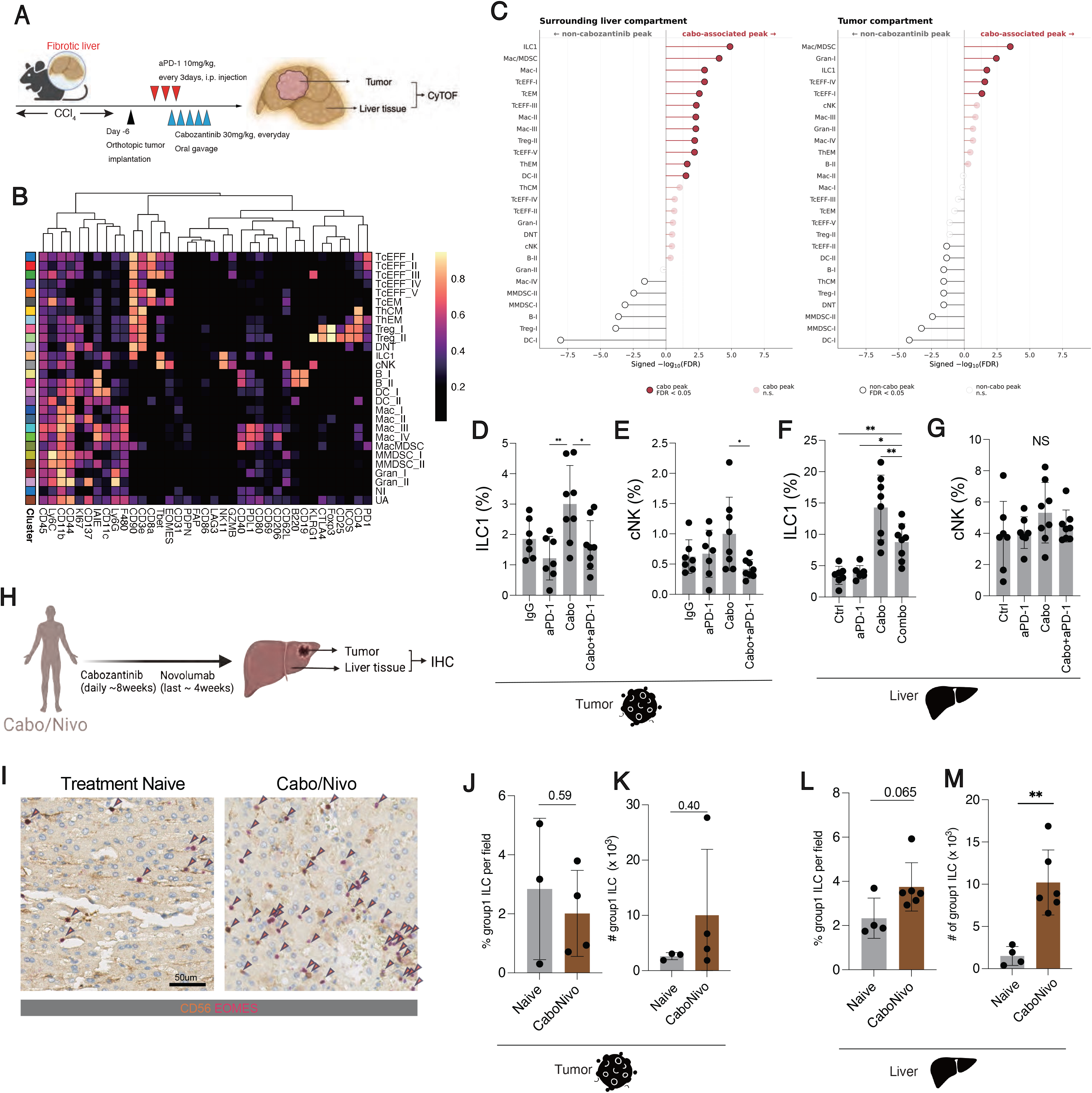
Cabozantinib/PD-1 blockade induces compartment-specific immune remodeling, with adaptive immune activation in tumors and group 1 innate lymphocyte expansion in fibrotic liver. (A) Experimental schema for CyTOF immune profiling of orthotopic RIL-175 tumors and matched surrounding fibrotic liver tissue after treatment with control, anti-PD-1 antibody, cabozantinib, or cabozantinib plus anti-PD-1. (B) Heatmap showing marker expression patterns across CyTOF-defined immune cell clusters. (C) Differential abundance analysis of immune clusters in surrounding liver and tumor compartments after treatment, highlighting non-cabozantinib-associated and cabozantinib-associated peaks. (D, E) Frequencies of ILC1 cells (D) and conventional NK cells (E) in tumor tissue. (F, G) Frequencies of ILC1 cells (F) and cNK cells (G) in surrounding liver tissue. (H) Schema of the neoadjuvant cabozantinib/nivolumab human HCC cohort used for tumor and non-tumor liver immunohistochemical analysis. (I) Representative CD56/EOMES duplex IHC images from treatment-naive and cabozantinib/nivolumab-treated liver tissue. Arrowheads indicate CD56-positive group 1 innate lymphocyte-enriched cells. Scale bar, 50 μm. (J-M) Quantification of CD56-positive group 1 innate lymphocyte-enriched populations in tumor tissue (J, K) and non-tumor liver tissue (L, M) from treatment-naive and cabozantinib/nivolumab-treated patients. Data are presented as mean +/- SD. *P < 0.05, **P < 0.01; NS, not significant.

In contrast, the surrounding fibrotic liver showed selective expansion of an NK-lineage cluster with low Eomes and high Tbet expression, consistent with an ILC1 phenotype (**Fig. S4H**), together with changes in effector T-cell and macrophage-associated clusters (**Fig. 3C, D, F** and **Fig. S4D-G**). A second NK cluster with features more consistent with circulating cNK cells was not comparably expanded (**Fig. 3E, G**). NKT-cell clusters were rare and did not substantially contribute in this model. These findings indicate that cabozantinib/ICB generates distinct immune outputs in tumor and liver: adaptive immune activation in tumor, but innate immune remodeling in the fibrotic liver (**Fig. 3A-G** and **Fig. S4A-H**).

To assess translational relevance, we analyzed liver and tumor specimens from HCC patients treated with neoadjuvant cabozantinib/nivolumab (**Fig. 3H-M**). Consistent with the murine data, treatment was associated with remodeling and expansion of CD56-positive group 1 innate lymphocyte-enriched populations in the liver, whereas changes in tumor tissue were less pronounced (**Fig. 3H-M**). Given the phenotypic complexity of human group 1 innate lymphocyte states and the limited marker set available in these specimens (17), these data are interpreted as human translational support for hepatic CD56-positive innate immune remodeling rather than definitive proof of expansion of a single ILC1 subset.

### NK1.1-positive innate lymphocytes mediate treatment-limiting hepatotoxicity without being required for tumor control

Given the selective expansion of NK-lineage populations in fibrotic liver, we next tested whether these cells functionally mediate hepatotoxicity (**Fig. 4A-L, Fig. S5A-E**). Tumor-bearing fibrotic mice were treated with NK1.1-depleting antibodies before cabozantinib/ICB (**Fig. S5A, B**). Depletion was confirmed in blood and liver by flow cytometry and histology (**Fig. S5C, D**). NK1.1-positive cell depletion significantly reduced treatment-associated weight loss, lowered liver enzyme elevations, decreased histologic liver injury, and preserved hepatocyte viability (**Fig. 4A, E-I**). Strikingly, this intervention restored survival benefit without impairing tumor regression (**Fig. 4B-D, Fig. S5E**). To complement these in vivo findings, we used an in vitro hepatic epithelial stress assay. When CCl_4_-damaged THLE-2 cells were co-cultured with human peripheral blood-derived NK-lineage cells, cabozantinib reduced the number of viable adherent THLE-2 cells and increased NK-cell attachment, consistent with enhanced susceptibility of stressed hepatic epithelial cells to NK-lineage cell–associated injury (**Fig. 4J–L**).

**Fig. 4.**
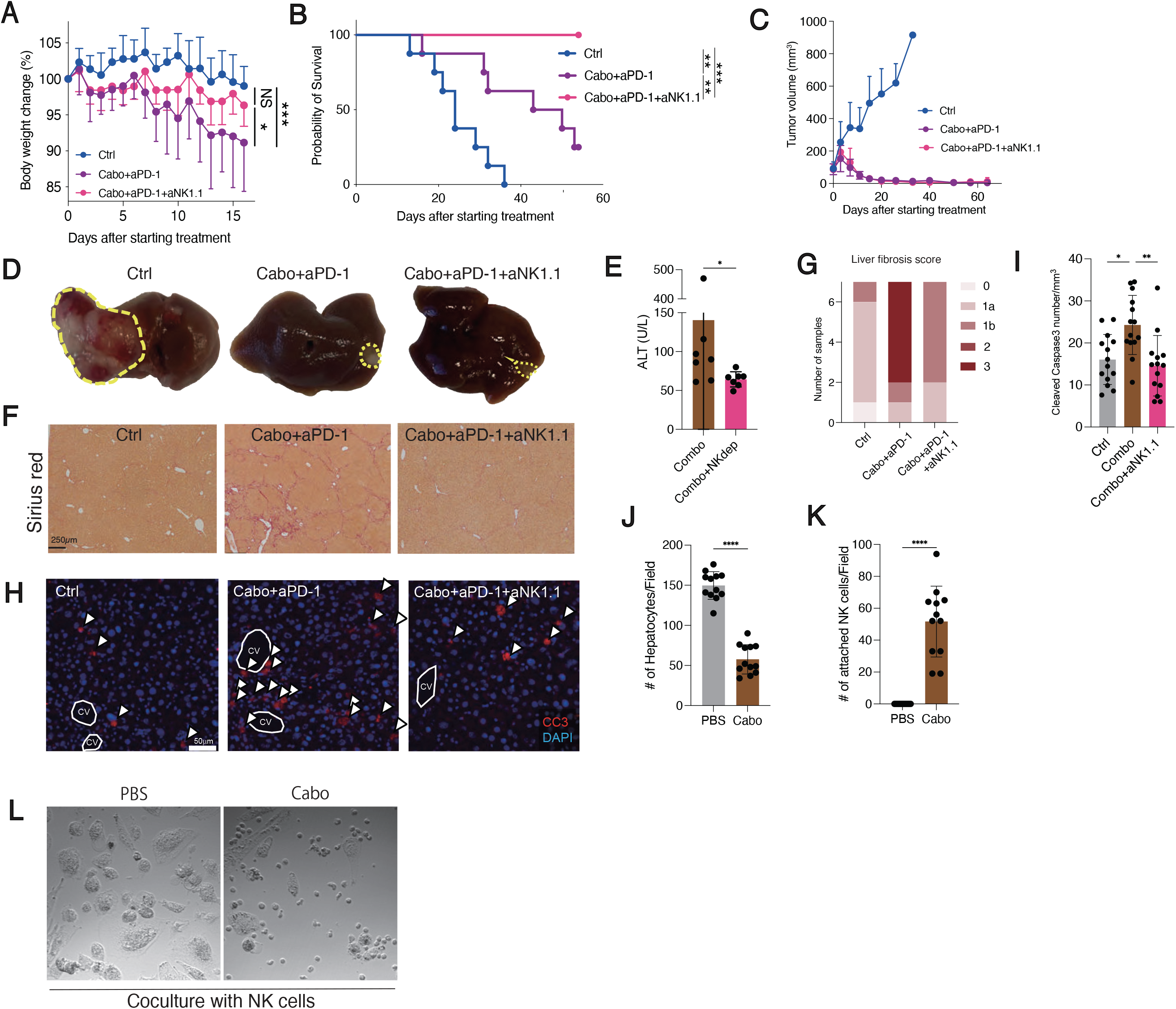
NK1.1-positive group 1 innate lymphocyte depletion mitigates hepatotoxicity and restores survival without impairing tumor control. (A-C) Body weight change (A), Kaplan-Meier survival curves (B), and tumor growth kinetics (C) in mice with CCl_4_-induced liver fibrosis and orthotopic RIL-175 HCC treated with control, cabozantinib plus anti-PD-1, or cabozantinib plus anti-PD-1 with anti-NK1.1-mediated group 1 innate lymphocyte depletion. (D) Representative macroscopic liver images from each treatment group. Dashed outlines indicate tumor areas. (E) Plasma ALT levels after cabozantinib/anti-PD-1 treatment with or without NK1.1-positive cell depletion. (F, G) Representative Sirius red staining (F) and liver fibrosis score distribution (G) in surrounding non-tumor liver tissue. Scale bar, 250 μm. (H, I) Representative cleaved caspase-3 immunofluorescence staining (H) and quantification of cleaved caspase-3-positive cells (I) in surrounding liver tissue. Arrowheads indicate cleaved caspase-3-positive cells; CV, central vein. Scale bar, 50 μm. (J-L) Quantification of viable THLE-2 hepatic epithelial cells (J), attached NK cells (K), and representative phase-contrast images (L) after co-culture of CCl_4_-damaged THLE-2 cells with peripheral blood-derived NK cells in the presence of PBS or cabozantinib. Data are presented as mean +/- SD. *P < 0.05, **P < 0.01, ***P < 0.001, ****P < 0.0001; NS, not significant.

These data identify the NK1.1-positive compartment as a key mediator of treatment-limiting liver injury while indicating that it is not required for the major antitumor effects of cabozantinib/ICB in this setting (**Fig. 4A-L**). Because anti-NK1.1 depletion targets a broader group 1 innate lymphocyte compartment rather than ILC1 exclusively, these results are interpreted as functional evidence for a pathogenic NK-lineage axis, with ILC1-like cells representing the leading candidate population based on the CyTOF and transcriptional data.

### Adaptive T cells do not mediate hepatotoxicity and may restrain innate immune-driven liver injury

To determine whether activated T cells contribute to liver damage, we depleted CD4-positive or CD8-positive T cells before cabozantinib/ICB treatment in fibrotic hosts without tumors (**Fig. S6A-J**). Neither CD4-positive nor CD8-positive T-cell depletion reduced treatment-associated toxicity (**Fig. S6B-D**). Instead, CD8-positive T-cell depletion worsened liver injury, increased fibrosis, and was associated with the expansion of group 1 innate lymphocytes in the liver (**Fig. S6E-H**). Conversely, NK1.1 depletion was associated with increased CD8-positive T-cell infiltration (**Fig. S6I, J**).

These reciprocal effects indicate that adaptive cytotoxic T cells are not the principal mediators of hepatotoxicity in this model and may instead constrain innate immune-driven liver injury (**Fig. S6A-J**). This differential effect also explains why antitumor efficacy is preserved after NK1.1 depletion despite a marked reduction in liver injury.

### Cabozantinib reprograms the fibrotic liver toward an innate immune-reactive stress state

To define molecular changes that might sensitize fibrotic livers to innate immune-mediated injury, we performed bulk RNA sequencing on liver tissues from mice with or without CCl_4_-induced fibrosis after cabozantinib treatment (**Fig. S7A-G**). Cabozantinib-treated fibrotic livers showed distinct transcriptional changes, including increased glycolytic and hypoxia-associated programs, enhanced fibroblast-related signatures, and reduced vascular programs, consistent with context-dependent hepatic stress (**Fig. S7B-E**). In parallel, ligands capable of engaging activating receptors on NK-lineage cells were altered in ways predicted to favor innate immune activation, including increased expression of *Raet1* and *Nectin2* and decreased expression of *H2T23* (**Fig. S7F**). Histologic examination of non-hepatic organs did not reveal comparable injury, supporting relative liver selectivity (**Fig. S7G**).

These data suggest that cabozantinib reprograms the fibrotic liver into a stress state that is permissive for innate immune recognition and injury (**Fig. S7A-G**). This effect was context-dependent and not a general consequence of therapy across tissues.

### Fibrosis and cabozantinib promote ILC1-like reprogramming with inflammatory effector programs

To define the cellular basis of innate immune remodeling, we performed single-cell RNA sequencing of liver populations after treatment (**Fig. 5A-H, Fig. S8A-H**). Clustering resolved hepatocytes, endothelial cells, T cells, and group 1 innate lymphocytes, which were further separated into cNK and ILC1-like states based on canonical transcriptional programs (**Fig. S8A-C**). Cabozantinib/ICB enriched the ILC1-like populations in fibrotic liver, consistent with the CyTOF and flow cytometric data (**Fig. 5A-C**). These cells showed increased proliferative activity and inflammatory activation, including enrichment of TNF-associated programs (**Fig. 5D-H, Fig. S8D-H**).

**Fig. 5.**
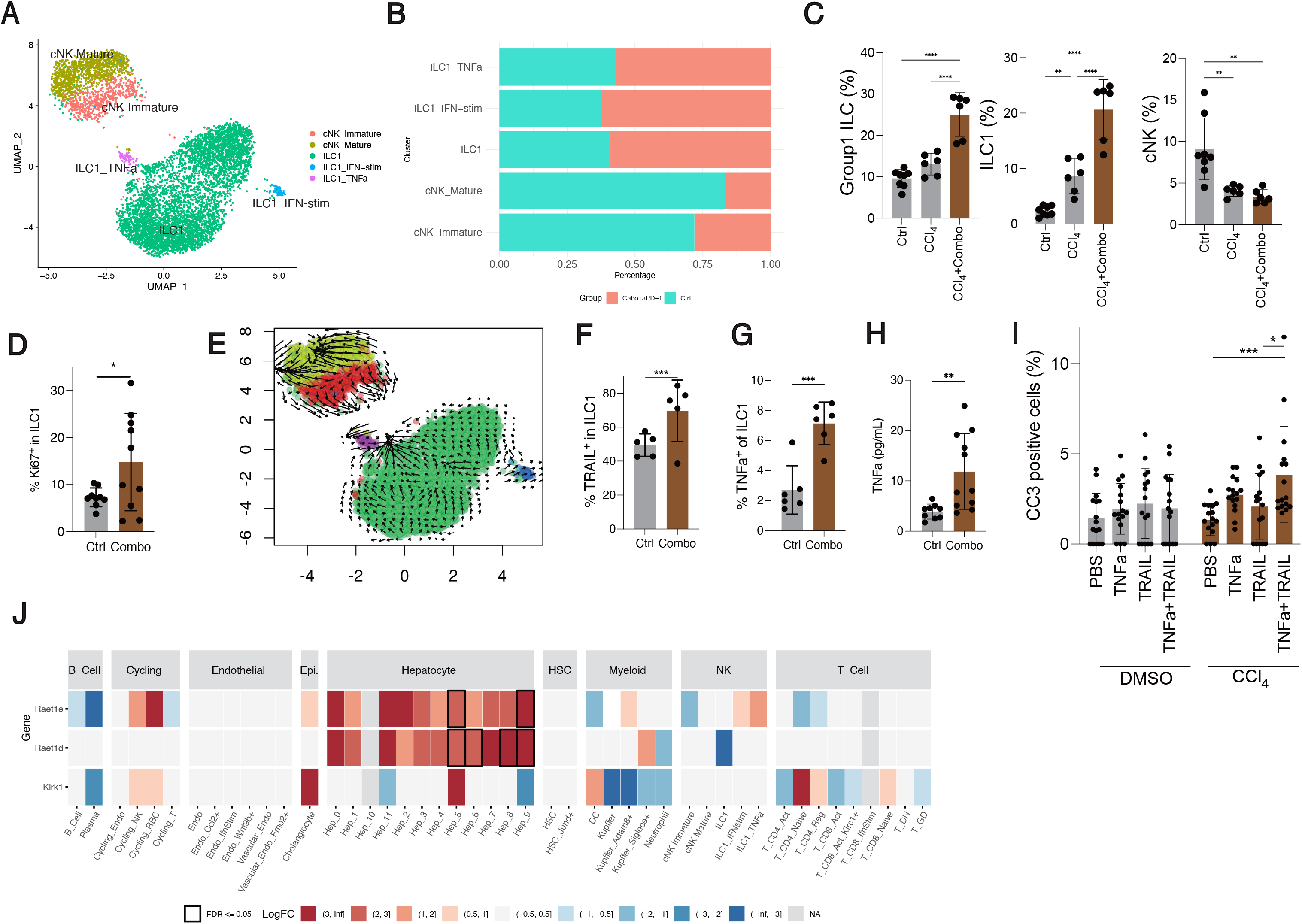
Cabozantinib/PD-1 blockade expands proliferative TNFα/TRAIL-associated ILC1-like cells and induces hepatocyte stress signaling in fibrotic liver. (A) UMAP visualization of scRNA-seq-defined group 1 innate lymphocyte subsets in fibrotic liver, including immature cNK cells, mature cNK cells, ILC1-like cells, interferon-stimulated ILC1-like cells, and TNFα-associated ILC1-like cells. (B) Relative proportions of group 1 innate lymphocyte subsets in control and cabozantinib/anti-PD-1-treated fibrotic liver. (C) Flow cytometric quantification of total group 1 innate lymphocytes, ILC1-like cells, and cNK cells in control liver, CCl_4_-induced fibrotic liver, and fibrotic liver treated with cabozantinib plus anti-PD-1. (D) Flow cytometric quantification of Ki67-positive ILC1-like cells, indicating increased proliferative activity after cabozantinib/anti-PD-1 treatment. (E) RNA velocity analysis showing directional cell-state dynamics toward the TNFα-associated ILC1-like state. (F, G) Flow cytometric quantification of TRAIL-positive ILC1-like cells (F) and TNFα-positive ILC1-like cells (G) after treatment. (H) Circulating TNFα levels after cabozantinib/anti-PD-1 treatment. (I) Quantification of cleaved caspase-3-positive cells after exposure of DMSO- or CCl_4_-pretreated THLE-2 hepatic epithelial cells to recombinant TNFα, TRAIL, or the combination. (J) Heatmap showing differential expression of selected NK-activating stress ligand and receptor-related genes across liver cell populations after cabozantinib/anti-PD-1 treatment. Outlined boxes indicate FDR <0.05. Data are presented as mean +/- SD. *P < 0.05, **P < 0.01, ***P < 0.001, ****P < 0.0001; NS, not significant.

Additional human NK-cell experiments showed that TGFβ, a fibrosis-associated cytokine, promoted partial acquisition of ILC1-like/tissue-resident features in CD56+ NK cells, including increased CD49a expression and expansion of EOMES-negative and TNFα-positive fractions. Separately, TNFα, particularly in combination with TRAIL, increased apoptosis of stressed hepatic epithelial cells, suggesting a potential effector mechanism linking inflammatory NK/ILC1-like programs to hepatotoxicity (**Fig.5I** and **Fig. S8I-M**). Together, these findings support fibrosis-associated reprogramming of NK-lineage cells toward an ILC1-like inflammatory state with hepatotoxic potential (**Fig. 5A-H and Fig. S8A-M**).

### Predicted hepatocyte-immune and endothelial-immune crosstalk is restricted to profiled cell populations

Single-cell analyses also identified candidate parenchymal–immune and endothelial–immune communication pathways. Cabozantinib/ICB induced hepatocyte stress-ligand programs and remodeled zonated endothelial compartments, with increased hypoxia-associated CAIX staining and altered endothelial Cxcl9/Cxcl16 expression (**Fig. 5J, Fig.S9A-H**) (18, 19). CellPhoneDB predicted CXCL9/CXCL10–CXCR3 and CXCL16–CXCR6 interactions between zonated endothelial cells, including peri-central endothelial cells, and ILC1-like populations (**Fig. S9I**). Because the single-cell dataset was generated using pre-enrichment and does not capture all hepatic immune populations, these interaction analyses were interpreted as candidate communication networks among profiled cell types only. Together, these data support a model in which hepatocyte stress and endothelial remodeling may contribute to NK-lineage retention or activation in the fibrotic liver microenvironment.

### Fibrotic liver promotes NK-to-ILC1-like conversion during cabozantinib/ICB

Because prior studies have shown that cNK cells can acquire ILC1-like properties in TGFβ-rich environments (12, 20, 21), we next tested whether the fibrotic liver niche promotes this transition during treatment (**Fig. 6A-D, Fig. S10A, B**). Receptor-ligand analyses identified interactions between group 1 innate lymphocytes and TGFβ-producing hepatic stromal populations (**Fig. 6A**). In adoptive-transfer experiments, GFP-positive cNK cells introduced into fibrotic recipients undergoing cabozantinib/ICB acquired a CD49a-positive CD49b-negative phenotype consistent with ILC1-like conversion (**Fig. 6B-D, Fig. S10A, B**).

**Fig. 6.**
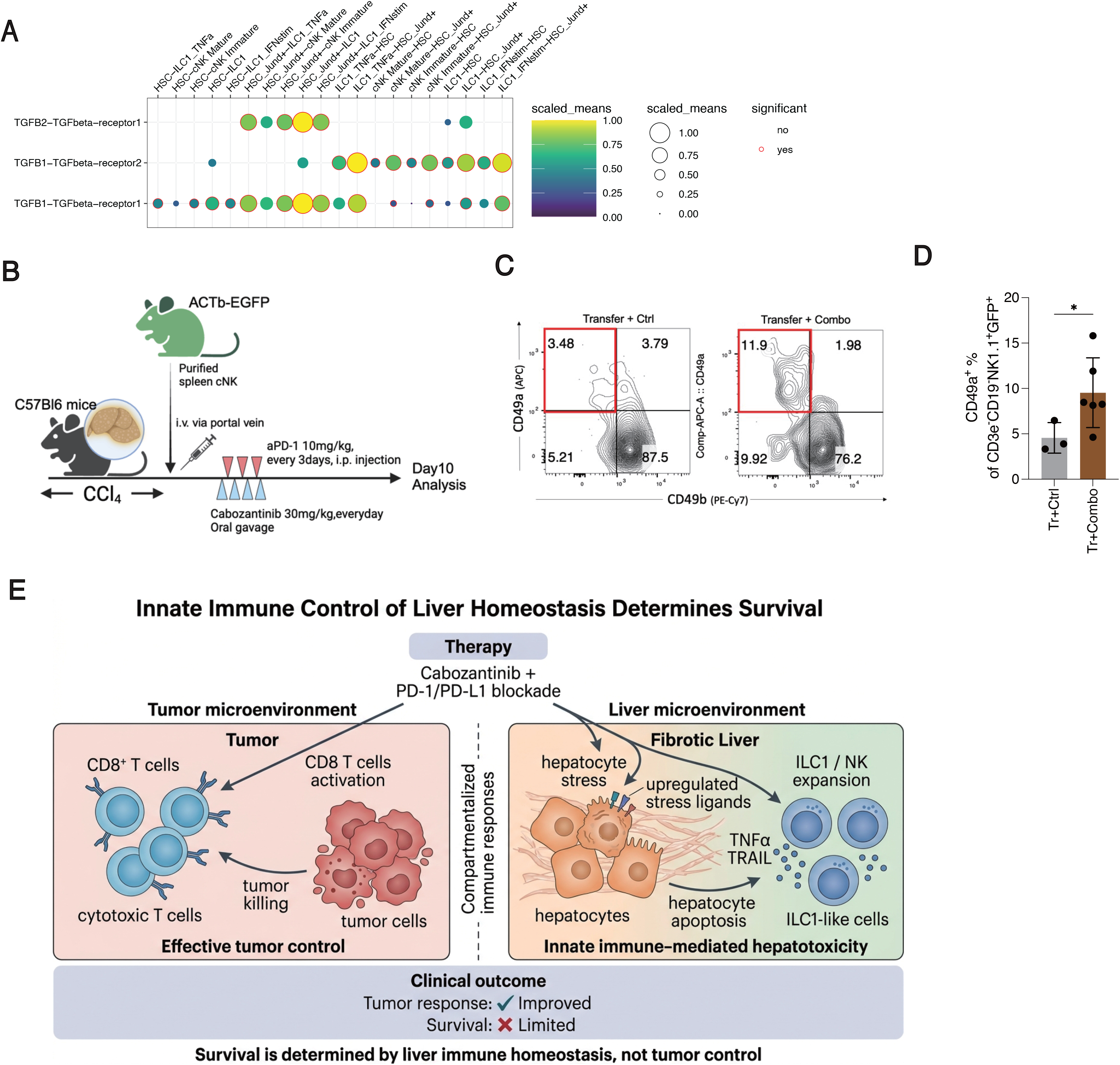
The fibrotic liver niche promotes NK-to-ILC1-like plasticity and innate immune-mediated hepatotoxicity. (A) Receptor-ligand interaction analysis showing predicted interactions between TGFβ ligands from hepatic stellate cell populations and TGFβ receptors on group 1 innate lymphocyte subsets. Dot size indicates scaled mean interaction strength; red outlines indicate statistically significant interactions. (B) Experimental schema for adoptive transfer of purified GFP-positive splenic cNK cells from ACTB-EGFP mice into CCl_4_-induced fibrotic recipient mice treated with control or cabozantinib plus anti-PD-1. (C) Representative flow cytometry plots showing CD49a and CD49b expression among donor-derived CD3e-negative CD19-negative NK1.1-positive GFP-positive cells after transfer. (D) Quantification of CD49a-positive cells among donor-derived CD3e-negative CD19-negative NK1.1-positive GFP-positive cells, indicating acquisition of an ILC1-like tissue-resident phenotype after cabozantinib/anti-PD-1 treatment. (E) Proposed model. Cabozantinib-based immunotherapy induces compartmentalized immune responses: cytotoxic CD8-positive T-cell activation mediates tumor control, whereas fibrotic liver promotes hepatocyte stress ligand expression, NK-lineage/ILC1-like expansion, TNFα/TRAIL signaling, hepatocyte apoptosis, and treatment-limiting hepatotoxicity. Thus, survival benefit is limited by liver immune homeostasis rather than tumor control alone. Data are presented as mean +/- SD. *P < 0.05.

These findings support a model in which fibrosis-associated cues, amplified by cabozantinib/ICB, promote phenotypic conversion of NK-lineage cells toward an ILC1-like state (**Fig. 6A-D**). This plasticity provides a plausible source of expanded pathogenic innate lymphocytes in the fibrotic liver during therapy.

### Integrated model of response-survival uncoupling in fibrotic liver

Collectively, these data support a model in which cabozantinib/ICB exerts spatially segregated and functionally divergent effects in HCC (**Fig. 6E**). Within tumors, treatment preserves or enhances cytotoxic immune programs and maintains antitumor efficacy (**Fig. 3C, Fig. S4B, C**). In the fibrotic liver, however, the same therapy promotes stress signaling, endothelial remodeling, and expansion or reprogramming of NK-lineage innate lymphocytes with ILC1-like features, resulting in hepatotoxicity that limits survival benefit (**Fig. 2E-H**, **Fig. 3C-M**, **Fig. 4A-I**, **Fig. 5A-J**, **Fig. 6A-D, Fig. S7-S10**). Thus, liver fibrosis uncouples tumor response from survival by narrowing the therapeutic index of MKI-ICB therapy through off-tumor innate immune-mediated liver injury (**Fig. 6E**).

## DISCUSSION

Survival after cabozantinib-based immunotherapy in HCC was determined in part by host liver condition rather than by tumor control alone. Across clinical analyses from COSMIC-312 and complementary murine models, baseline liver dysfunction was associated with inferior survival despite preserved tumor response, indicating that liver fibrosis can uncouple response from outcome in this treatment context (2,22). These findings support a host-integrated framework for therapeutic benefit in HCC, in which preservation of liver homeostasis is a parallel determinant of outcome alongside anticancer efficacy (2,3).

A key observation was that, in patients treated with cabozantinib plus atezolizumab, liver function stratified overall survival but not progression-free survival or tumor shrinkage. This pattern was not seen with sorafenib, in which liver status tracked more conventionally with both progression and survival. The distinction argues against a purely prognostic interpretation of liver reserve and instead supports a treatment-specific narrowing of the therapeutic index (2,22). In this model, patients with fibrotic or functionally compromised livers may still derive antitumor activity from cabozantinib/ICB, but that benefit is offset by reduced hepatic tolerance to treatment-associated injury (3,23).

The murine studies provide a mechanistic basis for this clinical dissociation. Across orthotopic, metastatic, and autochthonous HCC models, cabozantinib/ICB preserved tumor control irrespective of liver condition, yet survival benefit was lost selectively in fibrotic hosts. This loss of benefit was accompanied by body-weight loss, liver enzyme elevation, and worsening liver pathology, indicating that pre-existing fibrosis creates a vulnerable tissue state in which therapy-induced hepatotoxicity can override gains in tumor control. These data therefore support the interpretation that impaired survival in this setting reflects off-tumor liver injury rather than failure of anticancer efficacy.

Immune profiling further showed that cabozantinib/ICB induces spatially segregated immune effects. Within tumors, treatment enhanced cytotoxic CD8-positive T-cell programs, consistent with preserved antitumor immunity and with prior reports that cabozantinib can modulate the immune microenvironment (24−26). In contrast, the surrounding fibrotic liver exhibited expansion of NK-lineage populations with ILC1-like features. Data from patients treated with neoadjuvant cabozantinib nivolumab therapy support the translational relevance of this compartmentalized response by showing liver-predominant remodeling of CD56-positive group 1 innate lymphocyte-enriched populations after cabozantinib/nivolumab (13,27). Because the available human marker set does not permit definitive assignment of a single ILC1 subset, these findings are best interpreted as support for hepatic innate immune remodeling rather than direct proof of a one-to-one human correlate of the murine ILC1 population.

Functional depletion studies indicate that the NK1.1-positive compartment is a critical mediator of treatment-limiting hepatotoxicity. Removal of NK1.1-positive cells reduced liver injury and restored survival benefit without impairing tumor regression, whereas CD4-positive or CD8-positive T-cell depletion did not protect against toxicity. Indeed, CD8-positive T-cell depletion worsened liver injury, suggesting that adaptive cytotoxic T cells may restrain, rather than drive, the pathogenic innate response in fibrotic liver. These findings support a division of labor in which antitumor efficacy remains largely CD8-T-cell-associated, whereas treatment-limiting liver injury is disproportionately mediated by an NK-lineage compartment.

The mechanistic data further suggest that fibrosis and cabozantinib cooperate to reprogram the hepatic microenvironment toward innate immune-mediated injury. Bulk and single-cell transcriptomic analyses showed that cabozantinib-treated fibrotic livers acquire hypoxia- and stress-associated programs, altered vascular features, and increased expression of ligands capable of engaging activating receptors on group 1 innate lymphocytes. In parallel, innate lymphocyte states with ILC1-like features expanded and displayed inflammatory effector programs that included TNF-associated and TRAIL-associated signatures. Prior studies have shown that NK cells can acquire ILC1-like properties in TGF-β-rich settings (28−30), and our adoptive-transfer data support the idea that fibrosis-associated cues promote phenotypic conversion of conventional NK cells during therapy. Together, these observations are consistent with a model in which pre-existing fibrosis does not simply increase baseline inflammation but actively conditions the liver to respond to cabozantinib/ICB with pathogenic innate immune remodeling.

At the same time, anti-NK1.1 depletion targets a broader group 1 innate lymphocyte compartment, not ILC1 alone, and several innate populations can produce TNF and TRAIL under inflammatory conditions (31, 32). The data therefore support ILC1-like cells as major contributors within a pathogenic NK-lineage axis, rather than proving that they are the sole cellular source of hepatotoxic effector molecules.

These findings may also help explain a broader clinical problem in HCC. Several MKI-ICB combinations have shown meaningful antitumor activity yet failed to deliver proportional overall survival gains in patient populations with chronic liver disease (1,2). The present results suggest that, in HCC, therapeutic outcome may depend not only on whether a regimen activates antitumor immunity, but also on whether the injured liver can tolerate the accompanying host response (3,23). This concept of organ-specific vulnerability may be especially relevant in HCC, where the target organ is also the diseased organ and a major determinant of physiologic reserve.

The study also has practical translational implications. Readily available markers of liver condition, including APRI and ALBI, may help identify patients less likely to realize a survival benefit from cabozantinib-based combinations despite evidence of tumor response (33,34). More broadly, the data suggest that therapeutic strategies aimed at preserving liver homeostasis or modulating pathogenic innate immune activation could widen the therapeutic index of immunotherapy-based regimens in HCC. Whether such approaches should involve patient selection, toxicity-adapted dosing, or direct targeting of the implicated innate pathways will require prospective testing.

Our study has several limitations. First, the clinical analyses are retrospective and hypothesis-generating, although they are supported by orthogonal patient datasets and mechanistic animal studies. Second, the fibrosis models capture key aspects of chronic liver injury but do not fully reproduce the complexity of human cirrhosis and its systemic consequences (3,16). Third, the human neoadjuvant data provide translational support for hepatic innate immune remodeling, but they cannot establish direct NK-to-ILC1 conversion or definitive cellular causality in patients (13,27). Finally, the interventional data rely on broad NK1.1-based depletion, and more selective lineage-tracing or conditional targeting approaches will be needed to resolve the relative contributions of ILC1, cNK cells, and other minor NK1.1-positive populations.

Overall, these data support a model in which fibrosis-driven innate immune reprogramming narrows the therapeutic index of cabozantinib-based immunotherapy in HCC (**Fig. 6E**). In this framework, tumor control can be maintained while survival benefit is lost because treatment simultaneously activates an off-tumor hepatotoxic program in the fibrotic liver. By linking response-survival uncoupling to host liver homeostasis, this study provides a mechanistic basis for integrating liver condition more explicitly into therapeutic stratification and combination design for HCC.

## MATERIALS AND METHODS

### Study design

This study combined clinical data analyses, human translational specimen analyses, and mechanistic experiments in murine models to test whether host liver condition influences the relationship between tumor response and survival after cabozantinib-based immunotherapy in hepatocellular carcinoma (HCC). The work included retrospective analyses of clinical outcomes, pharmacovigilance and real-world safety datasets, orthotopic and autochthonous mouse HCC models with and without fibrosis, immune profiling, transcriptomic analyses, depletion studies, adoptive transfer experiments, and in vitro assays of hepatocyte injury.

### Clinical datasets and outcome definitions

Clinical analyses were performed using data from the randomized phase III COSMIC-312 trial in patients with previously untreated advanced HCC (2,22). Baseline liver function was assessed using the aspartate aminotransferase-to-platelet ratio index (APRI) and the albumin-bilirubin (ALBI) score. Outcomes analyzed included overall survival (OS), progression-free survival (PFS), objective tumor shrinkage, subsequent therapy use, and treatment-related liver toxicity. APRI was analyzed as both a dichotomous variable and a continuous variable. For dichotomized analyses, patients were grouped according to prespecified APRI thresholds. ALBI was analyzed alone and in combination with APRI using a validated composite threshold. Multivariable models adjusted for region, alpha-fetoprotein level, etiology, Eastern Cooperative Oncology Group (ECOG) performance status, and extrahepatic disease.

### Safety datasets

Cabozantinib-associated hepatotoxicity was further evaluated using adverse-event reports from the U.S. Food and Drug Administration pharmacovigilance database and an independent real-world cohort from the Massachusetts General Hospital Research Patient Data Registry. These datasets were used to evaluate hepatobiliary safety signals and liver enzyme abnormalities associated with cabozantinib-containing regimens.

### Human neoadjuvant specimens

Human liver and tumor specimens from patients with locally advanced HCC treated with neoadjuvant cabozantinib and nivolumab were analyzed to assess remodeling of the hepatic innate immune compartment (13). Liver and tumor samples were evaluated for CD56-positive group 1 innate lymphocyte-enriched populations using immunophenotyping and histologic approaches by a trained GI pathologist (JWL, Johns Hopkins University). These analyses were designed to provide translational support for the murine findings and to assess whether therapy-associated remodeling was detectable in human tissue.

### Cell lines and culture conditions

RIL-175 and HCA-1 murine HCC cells were maintained in Dulbecco’s modified Eagle medium supplemented with fetal bovine serum, pyruvate, and penicillin-streptomycin. Human THLE-2 hepatic epithelial cells were cultured under standard recommended conditions. Cell lines were authenticated before use and tested regularly for Mycoplasma contamination.

### Murine models

All animal studies were approved by the institutional animal care and use committee and performed according to institutional guidelines. Orthotopic HCC models were established by implantation of RIL-175 or HCA-1 cells into syngeneic recipient mice. To model chronic liver injury, mice received carbon tetrachloride (CCl4) before tumor implantation to induce fibrosis. This approach was used to create a fibrosis-primed host environment before initiation of anticancer therapy. An autochthonous fibrosis-associated HCC model was also used in Mst1-deficient; Mst2-floxed mice. These mice were exposed to the same fibrosis-inducing regimen and therapeutic interventions used in the orthotopic models.

### Treatment regimens

Mice bearing HCC received cabozantinib, immune checkpoint blockade, the combination, or control treatment according to the schedules specified for each experiment. Tumor burden was monitored longitudinally by the prespecified imaging or caliper-based methods appropriate to the model. Survival was recorded using protocol-defined humane endpoints. Body weight, serum alanine aminotransferase, and histopathology of non-tumor liver were used to assess treatment-related toxicity.

### Depletion studies

To evaluate the functional role of group 1 innate lymphocytes in treatment-associated liver injury, mice with fibrotic liver and established HCC were treated with anti-NK1.1 antibodies. These experiments targeted the broader NK1.1-positive compartment, including conventional NK cells, ILC1-like populations, and other NK1.1-expressing lymphocytes. Depletion efficiency was assessed by flow cytometry and tissue analysis. To assess the contribution of adaptive T cells, separate cohorts received anti-CD4 or anti-CD8 antibodies before cabozantinib-based therapy. These experiments were performed to determine whether CD4-positive or CD8-positive T cells were required for liver injury or protection.

### Histology and tissue analysis

Liver and tumor tissues were harvested at defined time points, fixed, embedded, and sectioned for histologic analysis. Standard stains and immunohistochemistry or immunofluorescence were used to evaluate fibrosis, hepatocyte injury, endothelial remodeling, and immune infiltration by a trained GI Pathologist (MMK, MGH). Non-tumor liver regions were analyzed separately from tumor tissue whenever possible.

### Isolation of liver and tumor immune cells

Liver and tumor tissues were mechanically and enzymatically dissociated to generate single-cell suspensions. Cells were enriched by density-gradient separation, washed, and stained for flow cytometry or downstream high-dimensional analyses. Fc receptor blocking was used before antibody staining. Viability, doublets, and dead cells were excluded during analysis.

### Flow cytometry

Flow cytometric analysis was performed on single-cell suspensions from liver, tumor, spleen, and blood as appropriate. Panels included markers to distinguish conventional NK cells, ILC1-like cells, T-cell subsets, and activation or effector markers such as Ki67, TNF, TRAIL, and lineage-associated transcription factors. Intracellular cytokine and transcription factor staining were performed after fixation and permeabilization using commercially available kits.

### Cytometry by time of flight

CyTOF was used to profile immune cells in matched tumor and non-tumor liver tissues. Samples were prepared using standard barcoding, surface staining, intracellular staining, and acquisition procedures (35, 36). After acquisition, normalization, debarcoding, dead-cell exclusion, and singlet gating were performed before downstream analysis. CyTOF data were clustered using unsupervised algorithms and visualized by dimensionality reduction methods. Cluster annotation was based on canonical marker expression and validated against complementary flow cytometry and histologic data. When technically feasible, selected absolute CD45-positive cell counts were calculated in addition to relative frequencies.

### Bulk RNA sequencing

Bulk RNA sequencing was performed on liver tissue collected from mice with and without CCl4-induced fibrosis after cabozantinib treatment. RNA was extracted, quality controlled, and used for library preparation and sequencing according to standard protocols. Differential expression analysis and gene set enrichment analysis were performed to define treatment-associated transcriptional programs, including hypoxia-related, glycolytic, fibroblast-associated, vascular, and stress-ligand signatures (37).

### Single-cell RNA sequencing

Single-cell RNA sequencing was performed on enriched liver immune populations after treatment to define immune states associated with cabozantinib-based therapy. Cells were processed using standard single-cell library preparation methods and sequenced on a high-throughput platform. Reads were aligned and processed to generate a gene-by-cell matrix for downstream analysis. Data were quality filtered, normalized, and analyzed using Seurat. Dimensionality reduction was performed using principal component analysis and UMAP, and batch effects were corrected with Harmony when needed. Cell clusters were annotated using canonical marker genes. Pseudobulk analyses were generated from raw counts for differential expression testing using edgeR (38).

### Receptor-ligand analysis and RNA velocity

Potential ligand-receptor interactions among profiled cell types were inferred using CellPhoneDB after ortholog mapping to human gene symbols (39). These analyses were used to identify candidate communication pathways among hepatocytes, endothelial cells, and group 1 innate lymphocyte populations. RNA velocity analysis was performed using velocyto and projected onto the UMAP embedding to assess directional cell-state transitions (40).

### Endothelial zonation analysis

Endothelial cells were analyzed for zonation-specific expression programs to assess periportal, mid-lobular, and pericentral identities. Zone scores were calculated from predefined marker gene sets using module scoring approaches (41, 42). These analyses were used to evaluate vascular remodeling under cabozantinib-based treatment.

### In vitro hepatocyte stress assays

THLE-2 cells were used to evaluate hepatic epithelial cell susceptibility to inflammatory injury. Cells were exposed to recombinant TNF, TRAIL, or both after stress conditioning when indicated. Apoptosis was quantified by cleaved caspase-3 staining and related viability readouts. These experiments were designed to test whether inflammatory mediators identified in vivo could directly induce hepatic epithelial cell injury.

### Coculture assays

Peripheral blood-derived NK-lineage cells were cocultured with THLE-2 cells in the presence or absence of cabozantinib. Cocultures were used to assess immune-cell interactions and hepatocyte susceptibility to NK-mediated cytotoxicity. In selected experiments, hepatic epithelial cell injury and cell-contact-dependent effects were quantified using microscopy and apoptosis-based readouts.

### Adoptive transfer experiments

To test whether conventional NK cells can acquire ILC1-like features in fibrotic liver, GFP-positive CD3-negative CD49a-negative CD49b-positive conventional NK cells were isolated from beta-actin GFP donor mice and transferred into wild-type recipient mice with established fibrosis. Recipients received cabozantinib-based therapy according to the experimental schedule. One week after transfer, recipient livers were analyzed by flow cytometry to determine acquisition of an ILC1-like phenotype by transferred cells.

### TGF-beta studies

TGF-beta pathway studies were performed to determine whether fibrotic liver microenvironmental signals contribute to conventional NK-to-ILC1-like reprogramming. These studies included measurement of TGF-beta expression in liver tissue, assessment of TGF-beta-producing stromal populations, and analyses of TGF-beta-associated signaling networks in the fibrotic liver. When performed, functional perturbation experiments evaluated whether TGF-beta-related cues influenced innate lymphocyte phenotype, proliferation, or hepatotoxic effector programs. Peripheral blood-derived NK-lineage cells were cultured with IL-15 in the presence or absence of TGFβ for 2 weeks. After culture, CD56-positive NK-lineage cells were analyzed for tissue-resident and ILC1-like markers by flow cytometry. After the induction, these NK-lineage cells were collected, and their tissue-resident markers were analyzed by flow cytometry.

### Statistics

Statistical analyses were performed using R and other software platforms as specified for each assay. Survival analyses used Kaplan-Meier methods, log-rank tests, Cox proportional hazards models, and competing-risk analyses where appropriate. Continuous associations were assessed using restricted cubic spline models. For experimental datasets, tests were selected based on the distribution and design of each comparison, and all tests were two-sided unless otherwise stated. Sample sizes, replicate numbers, and exact tests used for each figure are reported in the corresponding figure legends.

## Acknowledgments

We thank N.-J. Park, C Scheffold, M. Locker, H Ma, and D. Williamson (Exelixis) for providing COSMIC-312 data and analyses, and useful discussion. We are grateful to P. Ohashi (University of Toronto), H. Taniguchi, X. Liu, E. Krajnc, and M. Galvan (MGH) for valuable discussions, and M. Duquette, A. Khachatryan, C. Smith, and S. Roberge (MGH) for outstanding technical support.

## Funding

This work was supported by a sponsored research agreement for preclinical research from Exelixis (DGD). DGD’s research is supported by NIH grants R01CA260872, R01CA254351, and R01CA247441 and by Department of Defense PRCRP grant #W81XWH-19-1-0284. Mass cytometry work was also supported by P30CA006973 and S10OD034407 to WJH. S.M. received a postdoctoral fellowship from the Japan Society for the Promotion of Science and the MGH Fund for Medical Discovery (FMD) Fundamental Research Fellowship Award. The funders had no role in data analysis or the preparation of this article.

## Author contributions

conceptualization: SM, DGD, RKK, ML, MY, SD, RR; data curation: SM, HK, GB, AM, AM, TA, NJP, HM, CS, ML, RKK, WJH, RR; formal analysis: SM, HK, AM, AM, HM, SH, WJH; investigation: SM, HK, TA, GB, AM, TK, HM, SS, MMK, SD, RR, WJH; methodology: SM, HK, GB, AH, EMC, SMS, WJH, PH, RR; funding acquisition, project administration and supervision: DGD; validation: SM, TA, HK; visualization: SM, TA, HK, TK, WJH; writing – original draft: SM, DGD; writing – review & editing: all authors.

## Competing interests

D.G.D. received research grants from BMS, Bayer, and Surface Oncology. WJH reports patent royalties from Rodeo/Amgen, received research funding from Sanofi, NeoTX, Riboscience (to Johns Hopkins University), and speaking/travel honoraria from Exelixis and Standard BioTools. RKK reports research funding to institution from: Agios, Astra Zeneca, Bayer, BMS, Compass Therapeutics, Eli Lilly, EMD Serono, Exelixis, Genentech/Roche, Loxo Oncology, Merck, Novartis, Partner Therapeutics, QED, Relay Therapeutics, Servier, Surface Oncology, Taiho, Tyra Biosciences; consulting/advisory fees to institution from: Agios, Astra Zeneca, BMS, Exelixis, Ipsen, Merck; consulting/advisory fees to self from: Astra Zeneca, Compass, CVS Caremark, Elevar, GSK, Jazz, Moderna, Regeneron, Tyra Therapeutics, J-Pharma Inc.. The rest of the authors declare no competing interests.

## Data and materials availability

The authors declare that all data supporting the findings of this study are available within the paper and its supplementary material. Raw data are available upon reasonable request.

## Notes

### Competing Interest Statement

D.G.D. received consultant fees from Innocoll and research grants from BMS, Bayer, and Surface Oncology. No reagents from these companies were used in this study.

### Summary of Updates

This version has been substantially revised to clarify the central finding that liver fibrosis can uncouple tumor control from survival after cabozantinib based immunotherapy in hepatocellular carcinoma. The clinical analyses of COSMIC 312 were reorganized to emphasize the treatment specific relationship between baseline liver function, overall survival, progression free survival, tumor shrinkage, liver related toxicity, and subsequent therapy use. Additional APRI and ALBI based analyses and multivariable models were incorporated. The preclinical sections were revised to better align the mouse data with the clinical finding. Descriptions of the orthotopic, metastatic, and autochthonous HCC models were updated, including a clearer description of the Mst1 and Mst2 mouse experiments. The interpretation of hepatotoxicity, NK lineage cell depletion, CD4 and CD8 T cell depletion, hepatocyte stress signaling, and NK to ILC1 like plasticity was revised to avoid overstatement. New human translational analyses from neoadjuvant cabozantinib and nivolumab treated HCC samples were added to evaluate liver and tumor CD56 positive innate lymphocyte enriched populations. The terminology throughout the manuscript was standardized, using NK lineage cells, group 1 innate lymphocytes, and ILC1 like cells more precisely. Several figures, supplementary figures, figure legends, methods sections, and supplementary tables were updated for consistency. The revised manuscript also includes an expanded discussion of limitations, including the breadth of NK1.1 depletion, the restricted scope of cell interaction analyses, and the need for future lineage specific and human validation studies.

