## Supplemental figures and tables for "Liver fibrosis uncouples tumor control from survival after multikinase inhibitor immunotherapy in hepatocellular carcinoma"

#### Supplementary Figure Legends

**Supplementary Fig. S1. Baseline liver function stratifies survival and exploratory continuous APRI associations in COSMIC-312.** (A, B) Kaplan-Meier curves for progression-free survival (PFS) (A) and overall survival (OS) (B) in sorafenib-treated patients stratified by baseline APRI $<0.7$ versus APRI $>=0.7$. HRs, 95% CIs, median survival, P values, and numbers at risk are shown. (C, D) Exploratory restricted cubic spline models showing the association between baseline APRI as a continuous variable and hazard of progression (C) or death (D) in cabozantinib/atezolizumab-treated patients. Shaded areas indicate 95% confidence intervals. (E, F) Exploratory restricted cubic spline models showing the association between baseline APRI as a continuous variable and hazard of progression (E) or death (F) in sorafenib-treated patients. Shaded areas indicate 95% confidence intervals. (G, H) Kaplan-Meier curves for PFS (G) and OS (H) in cabozantinib/atezolizumab-treated patients stratified by combined APRI+ALBI score using a threshold of -2.46. HRs, 95% CIs, median survival, P values, and numbers at risk are shown.

**Supplementary Fig. S2. Cabozantinib-containing regimens are associated with liver-injury signals in pharmacovigilance and real-world datasets.** (A) Workflow for analysis of liver-related adverse events using the U.S. Food and Drug Administration adverse event reporting database, comparing cabozantinib with other multikinase inhibitors or bevacizumab. (B) Heatmap showing relative risks of reported liver-related adverse events, including immune-mediated hepatitis, increased liver function tests, drug-induced liver injury, autoimmune hepatitis, AST elevation, and ALT elevation, across systemic agents used in HCC. (C) Workflow for real-world analysis using the Partners/MGH database comparing cabozantinib-treated and sorafenib-treated patients after propensity score matching and descriptive statistical analysis.

**Supplementary Fig. S3. Cabozantinib/PD-1 blockade retains antitumor activity across HCC models but induces systemic toxicity in fibrotic hosts.** (A) Experimental schema for treatment of orthotopic HCC models in mice with normal liver. (B, C) Tumor growth kinetics (B) and Kaplan-Meier survival curves (C) in mice bearing orthotopic HCA-1 HCC in normal liver treated with control, anti-PD-1 antibody, cabozantinib, or cabozantinib plus anti-PD-1. (D) Experimental schema for treatment of orthotopic HCC models in mice with $CCl_4$-induced fibrotic liver. (E, F) Relative tumor size (E) and Ki67 index (F) in the autochthonous $Mst1^{-/-}Mst2^{f/-}$ HCC model with liver fibrosis after treatment. (G, H) Kaplan-Meier survival curves (G) and tumor growth kinetics (H) in mice bearing orthotopic HCA-1 HCC in fibrotic/damaged liver. (I, J) Body weight change in RIL-175 tumor-bearing mice with normal liver (I) or fibrotic/damaged liver (J) after treatment. (K) Body weight change in the autochthonous $Mst1^{-/-}Mst2^{f/-}$ HCC model with liver fibrosis after treatment. Data are presented as mean +/- SD. *P < 0.05, **P < 0.01, ***P < 0.001, ****P < 0.0001; NS, not significant.

**Supplementary Fig. S4. CyTOF profiling identifies compartment-specific immune remodeling after cabozantinib/PD-1 blockade.** (A) UMAP visualization of CyTOF-defined immune cell clusters from tumor and surrounding liver tissues. (B-G) Quantification of selected CyTOF-defined immune clusters after treatment. Tumor-associated immune changes included cytotoxic effector T-cell clusters (B, C), whereas surrounding liver changes included cytotoxic T-cell, macrophage, and NK-lineage clusters (D-G). (H) Marker expression profiles distinguishing ILC1 and cNK cell populations among CD3-negative NK1.1-positive cells. ILC1s were characterized by lower Eomes and CD62L expression, whereas cNK cells showed higher Eomes and CD62L expression. Data are presented as mean +/- SD. *P < 0.05, **P < 0.01, ***P < 0.001.

**Supplementary Fig. S5. Validation of NK1.1-positive cell depletion during cabozantinib/PD-1 blockade in fibrotic HCC-bearing mice.** (A) Experimental schema for NK1.1-positive cell depletion in mice with $CCl_4$-induced liver fibrosis and orthotopic RIL-175 HCC treated with cabozantinib plus anti-PD-1. Anti-NK1.1 treatment was initiated 1 day before cabozantinib/anti-PD-1 therapy. (B) Treatment-group allocation schema. (C, D) Representative flow cytometry plots (C) and histologic quantification (D) confirming depletion of NK1.1-positive cells after anti-NK1.1 treatment. (E) Individual tumor growth curves for control, cabozantinib plus anti-PD-1, and cabozantinib plus anti-PD-1 plus anti-NK1.1 treatment groups. Data are presented as mean +/- SD. ****P < 0.0001.

**Supplementary Fig. S6. CD4-positive or CD8-positive T-cell depletion does not mitigate cabozantinib/PD-1 blockade-induced hepatotoxicity in fibrotic liver.** (A) Experimental schema for CD4-positive or CD8-positive T-cell depletion in mice with $CCl_4$-induced liver fibrosis treated with cabozantinib plus anti-PD-1. Mice were sacrificed at day 18 after treatment initiation. (B) Body weight change after treatment with control, cabozantinib plus anti-PD-1, cabozantinib plus anti-PD-1 plus anti-CD8, or cabozantinib plus anti-PD-1 plus anti-CD4. (C, D) Serum AST (C) and ALT (D) levels after treatment. (E, F) Quantification of Sirius red-positive fibrotic area (E) and representative Sirius red staining (F) in liver tissue. Scale bar, 100 μm. (G, H) Quantification (G) and representative immunofluorescence images (H) of group 1 innate lymphocytes/NK1.1-positive cells in liver tissue. Scale bar, 50 μm. (I, J) Quantification (I) and representative immunofluorescence images (J) of CD8-positive cells after NK1.1-positive cell depletion. Arrowheads indicate positive cells. Data are presented as mean +/- SD. *P < 0.05, ***P < 0.001, ****P < 0.0001; NS, not significant.

**Supplementary Fig. S7. Cabozantinib induces hypoxia-associated transcriptional remodeling and NK-activating stress ligand programs in fibrotic liver.** (A) Experimental design for bulk RNA-seq analysis of healthy and $CCl_4$-induced fibrotic livers treated with control or cabozantinib. (B) Principal component analysis showing distinct transcriptional profiles according to liver fibrosis status and cabozantinib treatment. (C) Volcano plot showing differentially expressed genes in fibrotic liver after cabozantinib treatment. (D) Gene set enrichment analysis showing hallmark pathways enriched or depleted in fibrotic liver after cabozantinib treatment, including enrichment of hypoxia-associated pathways. (E) mMCP-counter analysis showing fibroblast-related and vessel-related transcriptional scores after cabozantinib treatment in fibrotic liver. (F) Expression of genes encoding NK-activating or inhibitory ligand-related molecules, including Raet1d, Raet1e, Nectin2, and H2t23, across treatment groups. (G) Representative histological images of colon and kidney tissue after treatment, showing no overt morphological injury outside the liver. Data are presented as mean +/- SD. *P < 0.05, **P < 0.01, ***P < 0.001.

**Supplementary Fig. S8. Single-cell transcriptomic and flow cytometric characterization of ILC1-like activation after cabozantinib/PD-1 blockade.** (A) Experimental workflow for scRNA-seq analysis of sorted liver cell populations from fibrotic mice treated with control or cabozantinib plus anti-PD-1. (B) Marker gene expression used to annotate major liver cell populations, including B cells, cholangiocytes, NK-lineage cells, endothelial cells, hepatocytes, hepatic stellate cells, myeloid cells, and T-cell subsets. (C) Expression of selected NK-lineage and ILC1-associated genes across cNK and ILC1 subsets. (D) Pathway enrichment analysis showing activation-associated programs in the TNFα-associated ILC1 subset, including TNFα signaling via NF-κB. (E) Representative flow cytometry plots showing TNFα expression among CD45-positive

CD3-negative NK1.1-positive CD49a-positive ILC1-like cells after treatment. (F-H) Flow cytometric quantification of granzyme B-positive ILC1 cells (F), IFNγ-positive ILC1 (G), and granzyme C-positive ILC1 (H). (I) Quantification of cleaved caspase-3-positive cells after exposure of THLE-2 hepatic epithelial cells to increasing concentrations of recombinant TNFα. (J) Experimental design for the phenotype of peripheral blood-derived NK cells from healthy donors in the presence of DMSO or TGFβ. (K-M) Flow cytometric quantification of CD49a-positive cells (K), EOMES-negative cells (L), and TNFα-positive cells (M) in CD56-positive cells. The plots were replicated from a single donor, and 2 donors were analyzed independently. Data are presented as mean +/- SD. *$P < 0.05$, **$P < 0.01$, ***$P < 0.001$; NS, not significant.

**Supplementary Fig. S9. Cabozantinib/PD-1 blockade induces vascular rarefaction, hypoxia, and endothelial chemokine remodeling in fibrotic liver.** (A) Flow cytometric quantification of CD31-positive CD45-negative endothelial cells in liver tissue after control or cabozantinib/anti-PD-1 treatment. (B, C) Representative immunofluorescence images of CA-IX staining (B) and quantification of CA-IX-positive liver area (C) after treatment. Scale bar as shown. (D) UMAP visualization of scRNA-seq-defined liver endothelial cell subsets annotated by zonation signatures, including periportal, mid-lobular, and peri-central endothelial populations. (E) UMAP visualization of endothelial cell subsets by treatment group. (F) Relative proportions of endothelial cell subsets in control and cabozantinib/anti-PD-1-treated liver. (G) Expression of Cxcl9 and Cxcl16 across endothelial zonation identities. (H) Expression of Cxcl9 and Cxcl16 in endothelial cells from control and cabozantinib/anti-PD-1-treated liver. (I) Receptor-ligand interaction analysis showing predicted chemokine interactions between endothelial zonation subsets and group 1 innate lymphocyte subsets, including CXCL9-CXCR3, CXCL10-CXCR3, and CXCL16-

CXCR6 axes. Dot size indicates scaled mean interaction strength; red outlines indicate statistically significant interactions. Data are presented as mean +/- SD. *P < 0.05, ****P < 0.0001.

**Supplementary Fig. S10. Characterization of transferred cNK cells and expansion of NK1.1-positive cells after cabozantinib/PD-1 blockade.** (A) Representative flow cytometry plot showing CD49a and CD49b expression among splenic live CD45-positive CD3-negative CD19-negative NK1.1-positive lymphocytes used for adoptive transfer, confirming enrichment of CD49a-negative CD49b-positive cNK cells. (B) Quantification of NK1.1-positive cells among CD3e-negative CD19-negative lymphocytes in recipient mice after adoptive transfer and treatment with control or cabozantinib plus anti-PD-1. Data are presented as mean +/- SD. **P < 0.01.

#### Supplementary Table Legends

**Supplementary Table S1. Multivariable Cox regression analysis of overall survival according to baseline clinical variables.**

Multivariable Cox proportional hazards models evaluating the association between baseline clinical variables and overall survival in patients with advanced HCC treated with cabozantinib plus atezolizumab. Liver function was modeled using either APRI <0.7 or the combined ALBI+APRI score < -2.46, together with region, etiology, extrahepatic disease, AFP level, and ECOG performance status. Hazard ratios (HRs), 95% confidence intervals (CIs), and P values are shown. AFP, alpha-fetoprotein; ALBI, albumin-bilirubin index; APRI, aspartate aminotransferase-to-platelet ratio index; CI, confidence interval; ECOG, Eastern Cooperative Oncology Group; HBV, hepatitis B virus; HCV, hepatitis C virus; HCC, hepatocellular carcinoma; HR, hazard ratio.

**Supplementary Table S2. Baseline demographics and clinical characteristics of patients enrolled in COSMIC-312.** Baseline demographics, tumor characteristics, liver function parameters, and disease characteristics of patients with advanced HCC treated with cabozantinib plus atezolizumab, sorafenib, or single-agent cabozantinib in COSMIC-312. Data are presented as median [interquartile range] or number of patients (%). ALBI, albumin-bilirubin index; BCLC, Barcelona Clinic Liver Cancer; ECOG, Eastern Cooperative Oncology Group; HBV, hepatitis B virus; HCV, hepatitis C virus; HCC, hepatocellular carcinoma; IQR, interquartile range.

**Supplementary Table S3. Post-protocol therapy use and time to first systemic therapy after study drug discontinuation.** Summary of post-protocol systemic, local, and unknown therapies among patients treated with cabozantinib plus atezolizumab, single-agent cabozantinib, or sorafenib. Time to first post-protocol systemic therapy is shown as median, range, and 25th-75th percentile among patients who received subsequent systemic therapy. Cabo, cabozantinib.

**Supplementary Table S4. Baseline demographics and liver enzyme profiles of cabozantinib- or sorafenib-treated patients after propensity score matching.** Baseline demographic characteristics and liver enzyme abnormalities in a real-world cohort of patients treated with cabozantinib or sorafenib after propensity score matching. AST and ALT grades, as well as the frequencies of AST and ALT elevation, are shown for the overall cohort and by treatment group. P values compare cabozantinib-treated and sorafenib-treated patients. ALT, alanine aminotransferase; AST, aspartate aminotransferase; IQR, interquartile range.

**Supplementary Table S5. Association between baseline clinical factors and cabozantinib-induced liver injury.** Logistic regression analysis evaluating baseline clinical factors associated with cabozantinib-induced liver injury. Odds ratios (ORs), 95% confidence intervals (CIs), and P values are shown for race, age, sex, ALBI, ethnicity, and APRI. ALBI, albumin-bilirubin index; APRI, aspartate aminotransferase-to-platelet ratio index; AST, aspartate aminotransferase; CI, confidence interval; OR, odds ratio.

**Supplementary Table S6. Summary of cabozantinib-associated hepatotoxicity reported in clinical trials.** Summary of AST and ALT elevations reported in representative clinical trials of cabozantinib or cabozantinib-based immune checkpoint blockade compared with comparator arms. The table includes COSMIC-312, COSMIC-021, CELESTIAL, and NCT00940225. ALT, alanine aminotransferase; AST, aspartate aminotransferase; HCC, hepatocellular carcinoma; ICB, immune checkpoint blockade; N/A, not available; RCC, renal cell carcinoma.

**Table R1. Antibodies used for CyTOF immune profiling.** The table lists the metal isotope channel, target antigen, antibody clone, whether the antibody was custom-conjugated, and commercial source. "X" indicates custom-conjugated antibodies.

Oncology, Merck, Novartis, Partner Therapeutics, QED, Relay Therapeutics, Servier, Surface Oncology, Taiho, Tyra Biosciences; consulting/advisory fees to institution from: Agios, Astra Zeneca, BMS, Exelixis, Ipsen, Merck; consulting/advisory fees to self from: Astra Zeneca, Compass, CVS Caremark, Elevar, GSK, Jazz, Moderna, Regeneron, Tyra Therapeutics, J-Pharma Inc.. The rest of the authors declare no competing interests.

**Data and materials availability:** The authors declare that all data supporting the findings of this study are available within the paper and its supplementary material. Raw data are available upon reasonable request.

### Supplementary Fig.S1

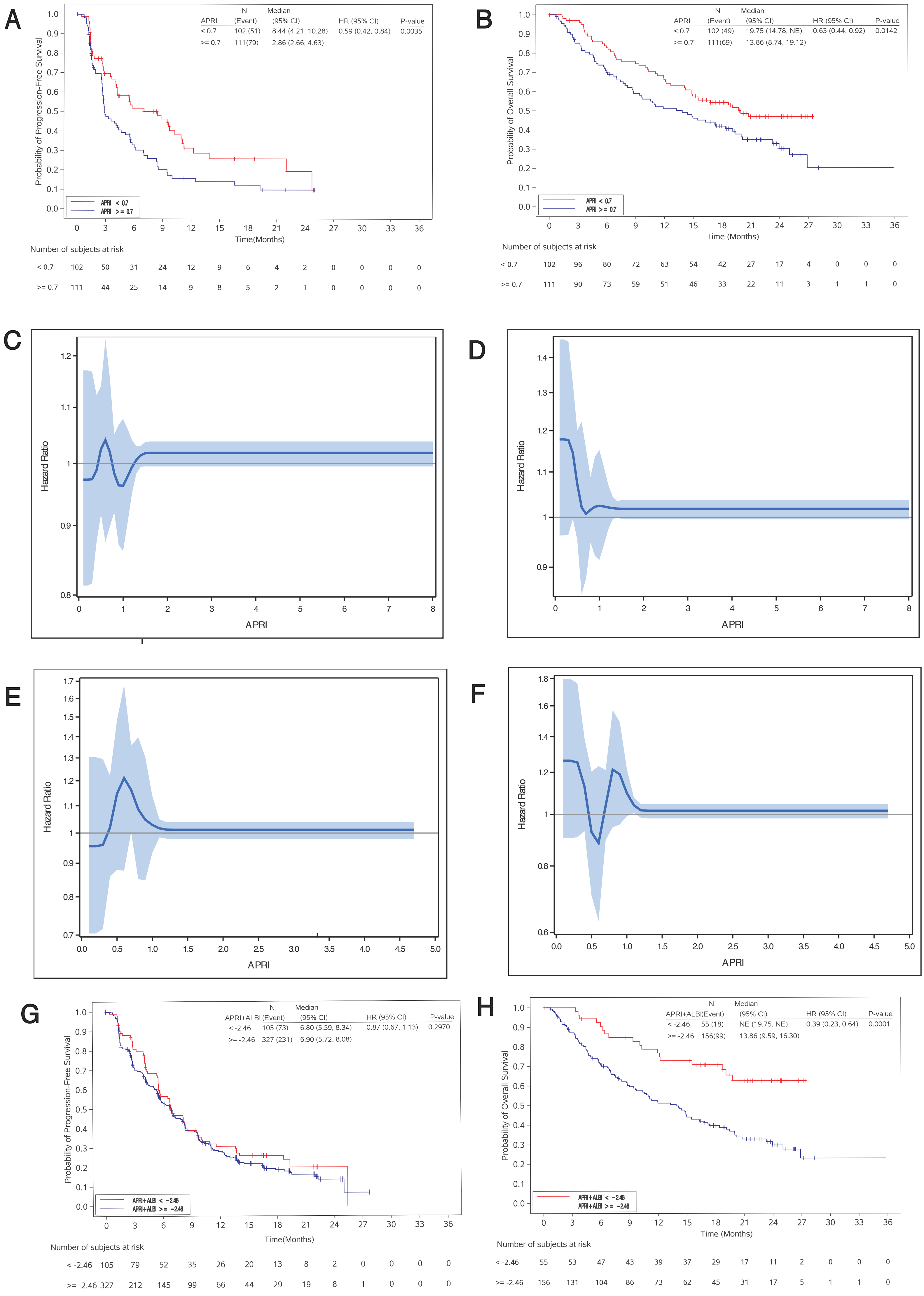

Supplementary Fig.S2

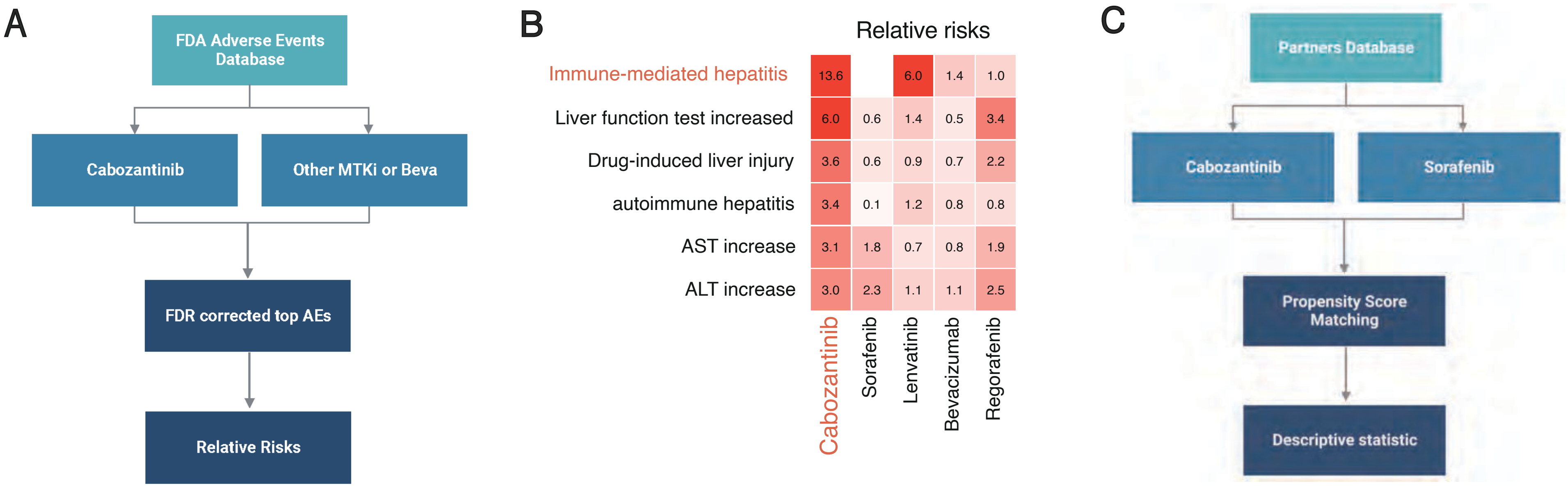

#### Supplementary Fig.S3

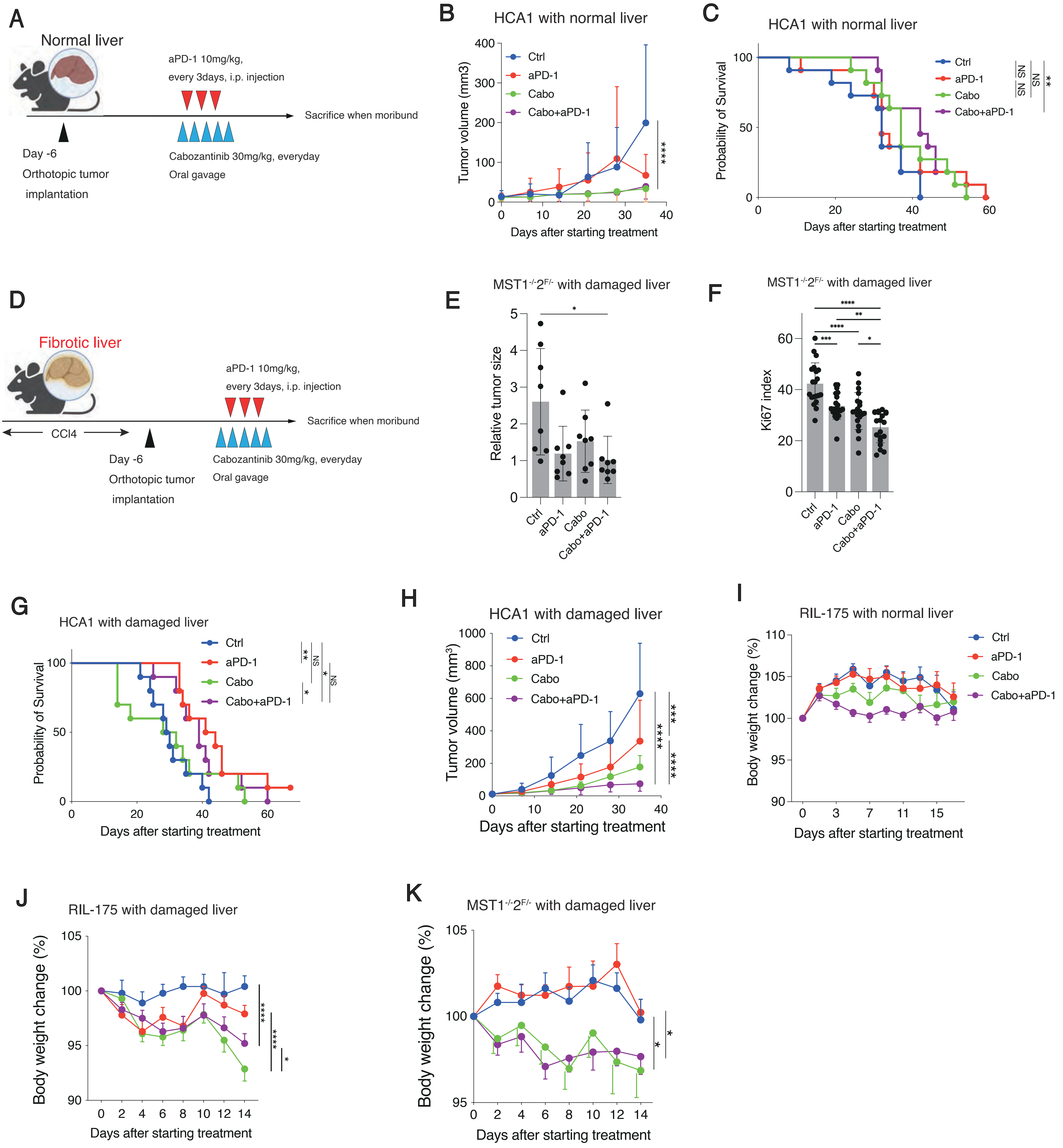

### Supplementary Fig.S4

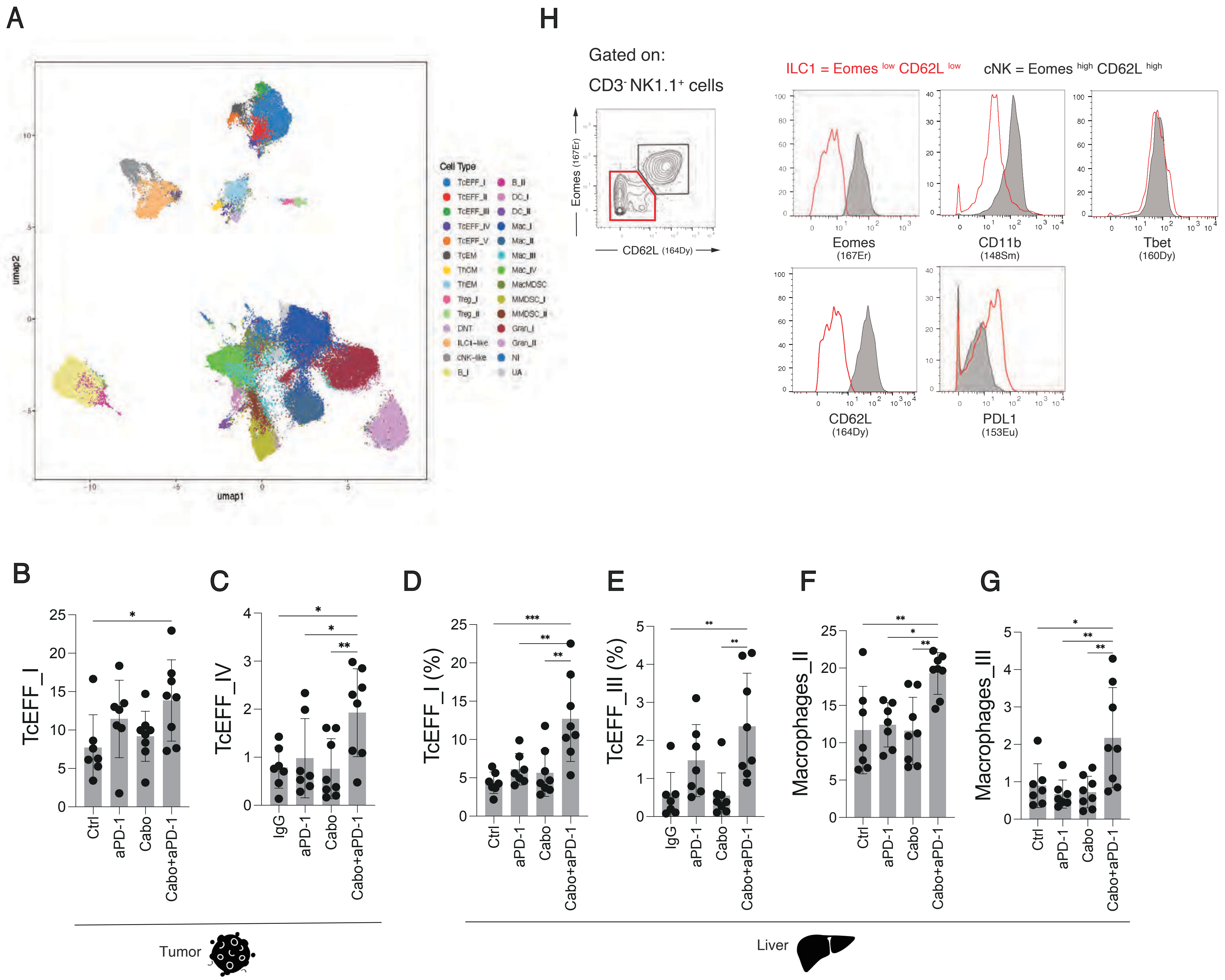

### Supplementary Fig.S5

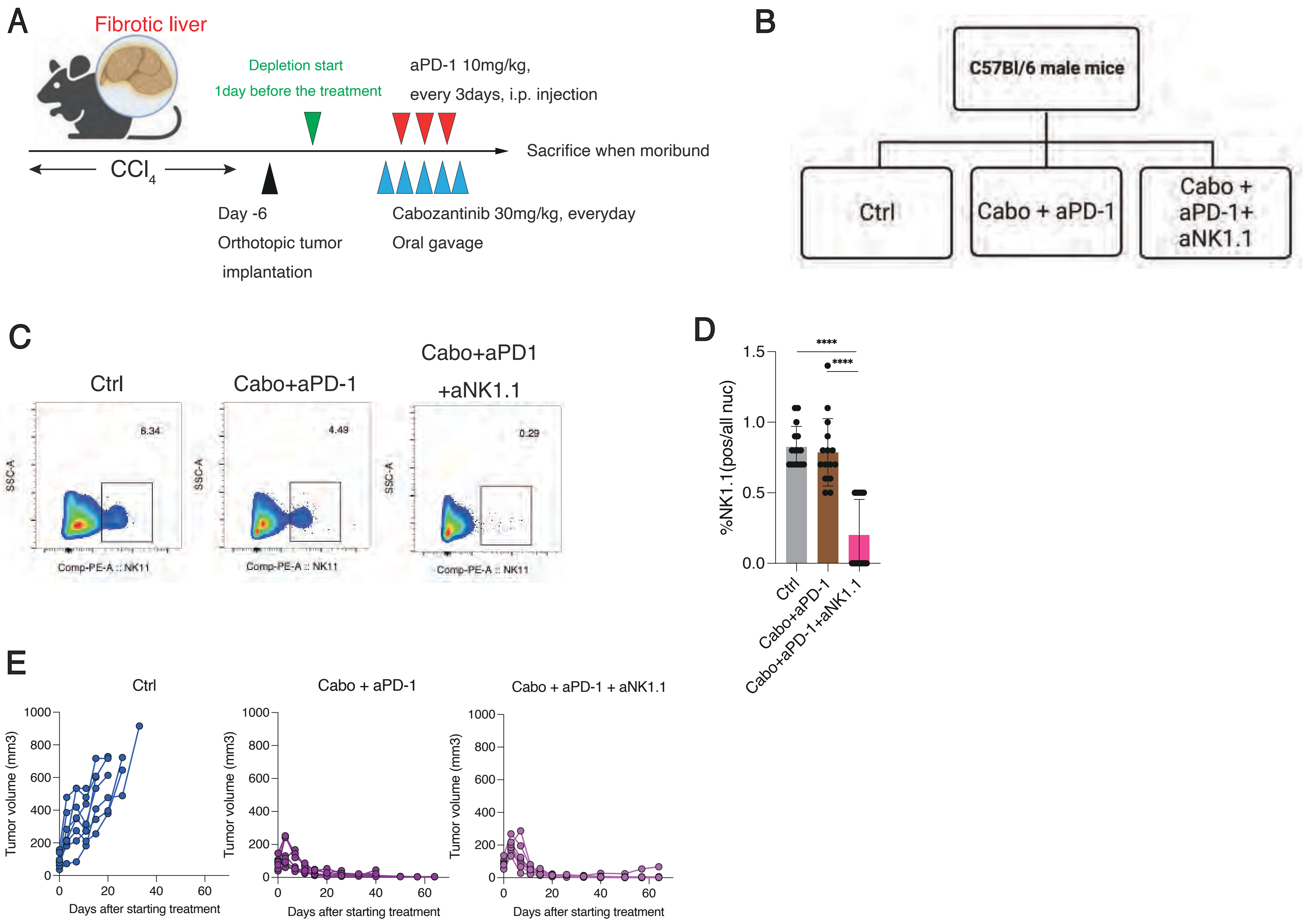

### Supplementary Fig.S6

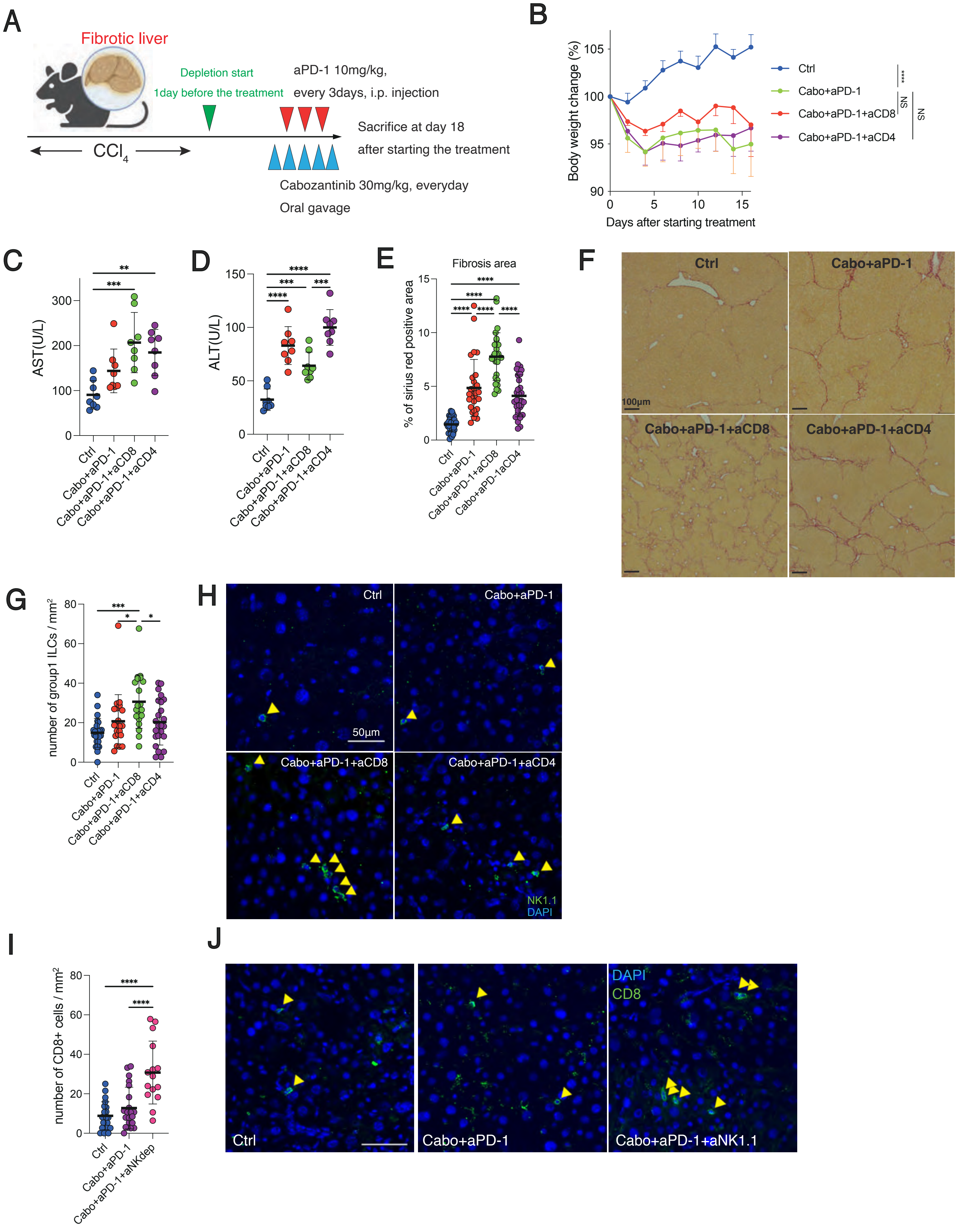

#### Supplementary Fig.S7

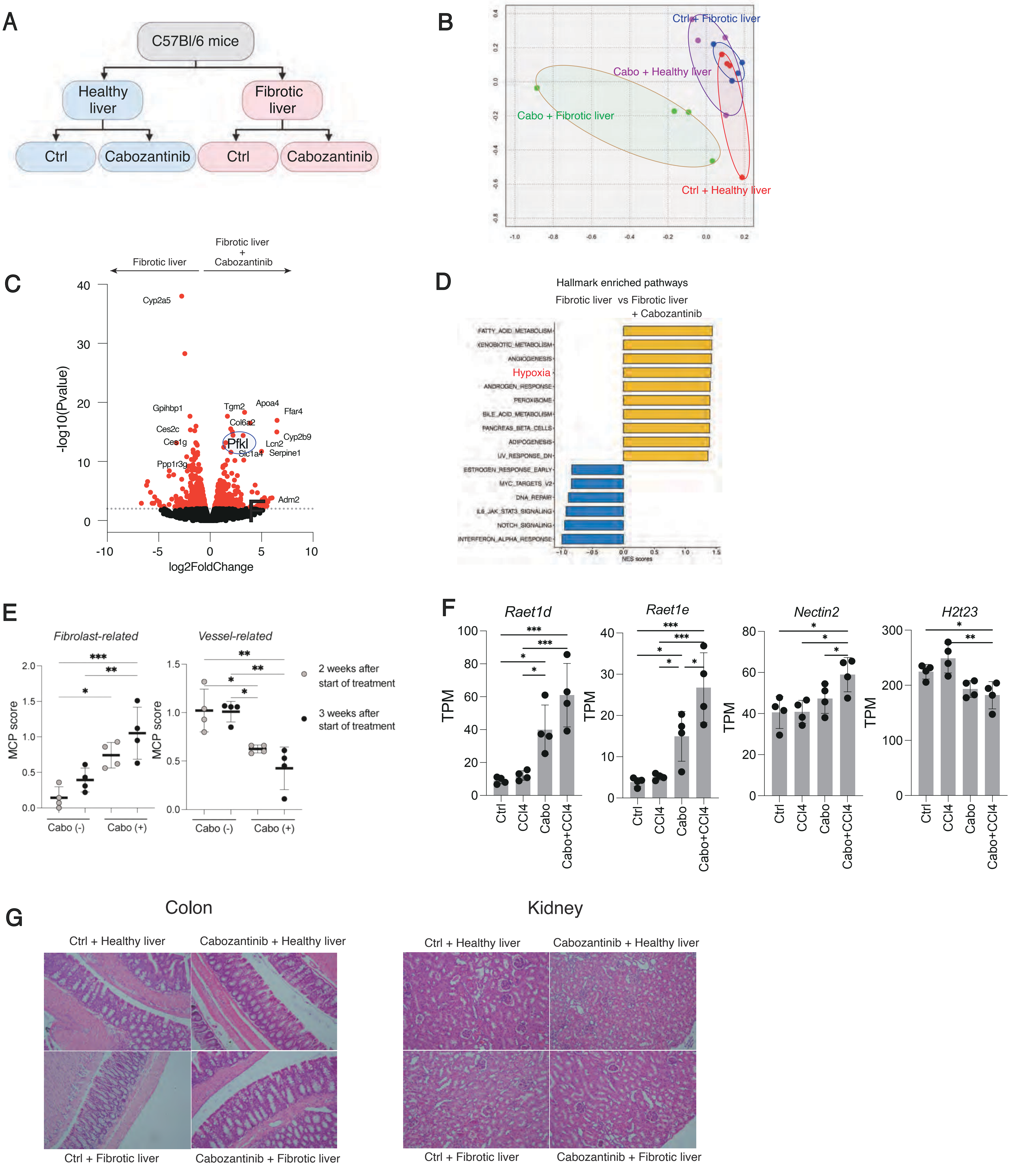

### Supplementary Fig.S8

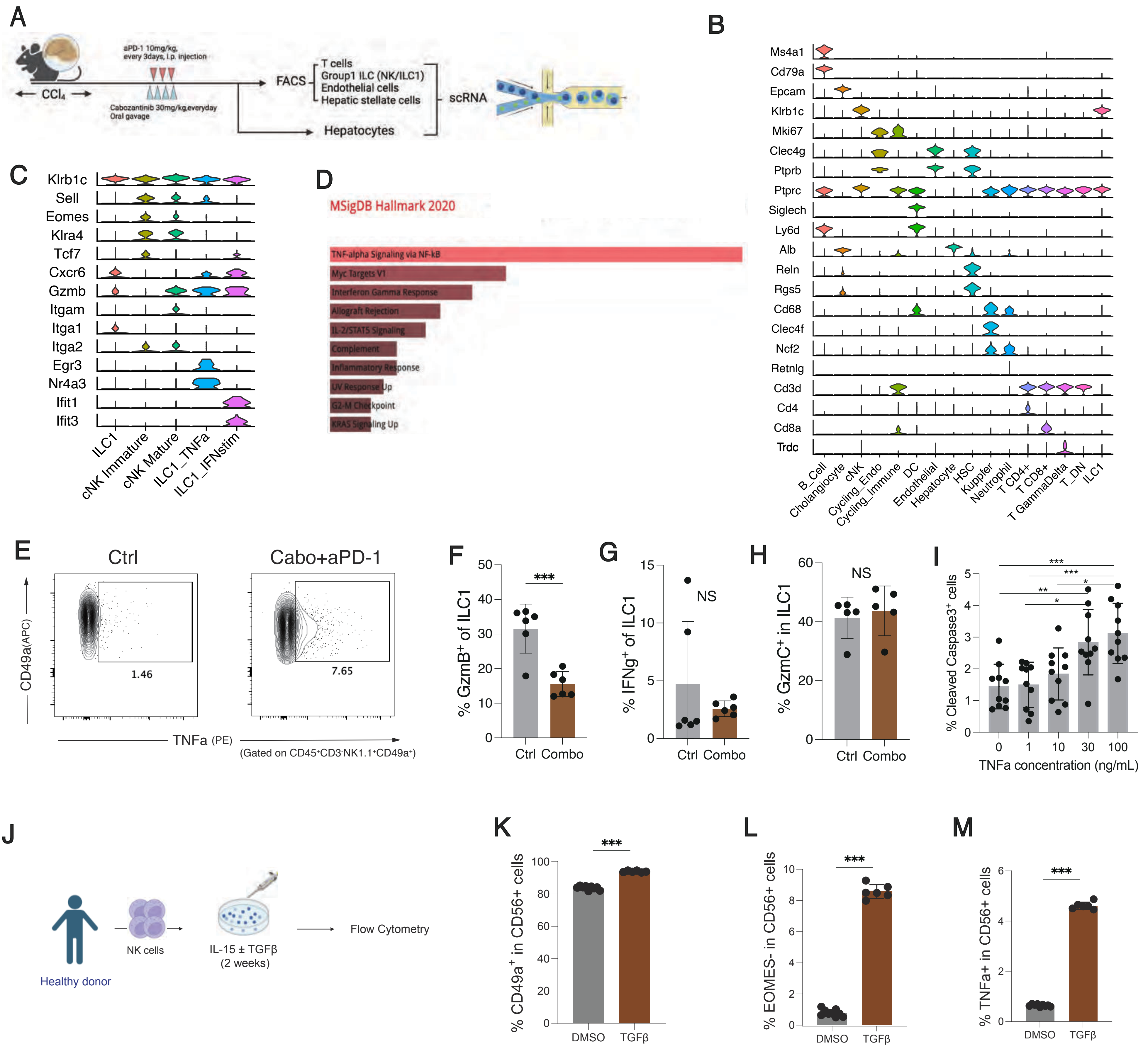

#### Supplementary Fig.S10

A

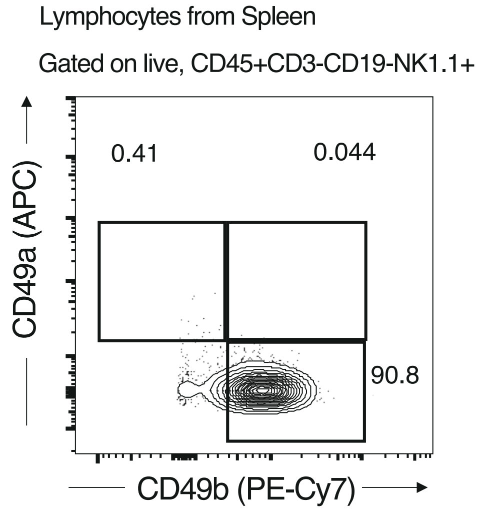


B

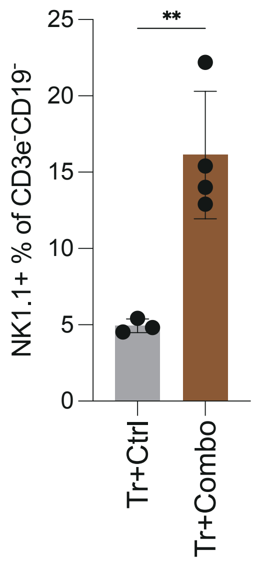

### Supplementary Fig. S9

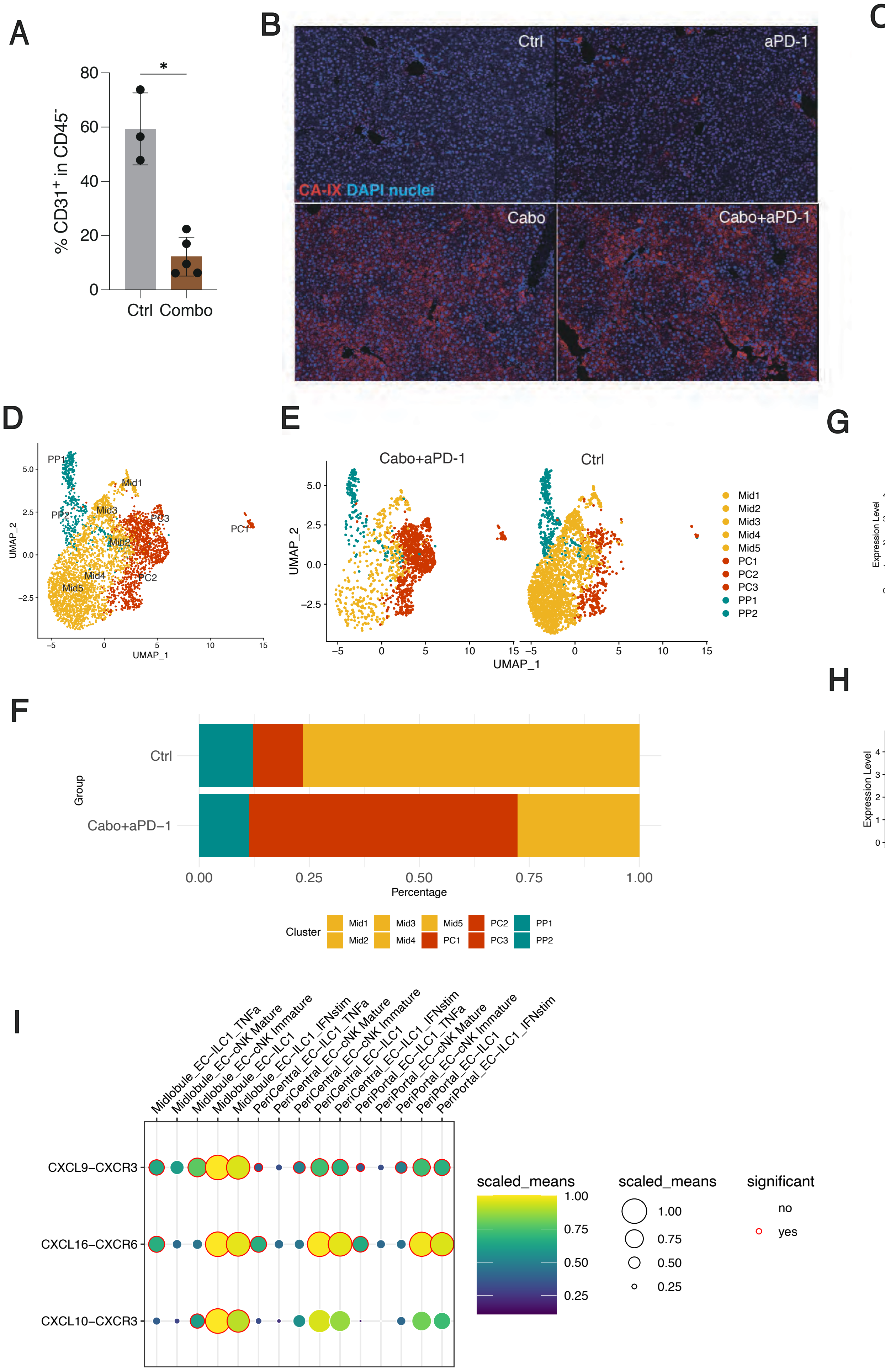

### Supplemental Table 1

**Multivariable Cox regression analysis for overall survival with baseline clinical variables.**

| Variable | HR (95% CI) | p-value | Variable | HR (95% CI) | p-value |
|---|---|---|---|---|---|
| Region: Asia | 0.88 (0.62–1.24) | 0.463 | Region: Asia | 0.84 (0.59–1.18) | 0.311 |
| Etiology: HBV ± HCV | 0.83 (0.57–1.19) | 0.3 | Etiology: HBV ± HCV | 0.93 (0.64–1.35) | 0.7 |
| Etiology: HCV only | 1.00 (0.75–1.34) | 0.995 | Etiology: HCV only | 1.02 (0.76–1.35) | 0.918 |
| Extrahepatic disease: Absent | 0.89 (0.68–1.16) | 0.389 | Extrahepatic disease: Absent | 0.87 (0.66–1.14) | 0.321 |
| AFP < 400 ng/mL | 0.71 (0.55–0.92) | **0.009** | AFP < 400 ng/mL | 0.69 (0.52–0.89) | **0.004** |
| ECOG 0 | 0.68 (0.53–0.88) | **0.003** | ECOG 0 | 0.67 (0.52–0.86) | **0.002** |
| APRI < 0.7 | 0.73 (0.57–0.95) | **0.017** | ALBI+APRI < –2.46 | 0.47 (0.34–0.65) | **<0.001** |

Abbreviations: AFP, alpha-fetoprotein; ECOG, Eastern Cooperative Oncology Group; ALBI, albumin–bilirubin index; APRI, AST to platelet ratio index.

### Supplemental Table 2

#### Baseline demographics and clinical characteristics of the study population

| Characteristic | Cab+Atezo (n=432) | Sorafenib (n=217) | Cabo (n=188) |
|---|---|---|---|
| Age, years (median [IQR]) | 64 [58–70] | 64 [57–71] | 64 [57–71] |
| Sex | | | |
| Male | 306 (71%) | 186 (86%) | 158 (84%) |
| Female | 126 (29%) | 30 (14%) | 30 (16%) |
| Race† | | | |
| White | 213 (52%) | 113 (52%) | 91 (48%) |
| Asian | 151 (35%) | 63 (29%) | 58 (31%) |
| Black or African American | 11 (3%) | 8 (4%) | 6 (3%) |
| Other/Not reported | 57 (13%) | 33 (15%) | 33 (18%) |
| Geographical region | | | |
| Asia | 210 (49%) | 63 (29%) | 58 (31%) |
| Other | 222 (51%) | 154 (71%) | 130 (69%) |
| Etiology | | | |
| HBV (±HCV) | 127 (29%) | 64 (29%) | 59 (31%) |
| HCV (no HBV) | 119 (28%) | 68 (31%) | 61 (32%) |
| Non-viral | 186 (43%) | 85 (39%) | 68 (36%) |
| ECOG performance status | | | |
| 0 | 206 (48%) | 107 (49%) | 96 (51%) |
| 1 | 226 (52%) | 110 (51%) | 92 (49%) |
| Extrahepatic disease | 292 (68%) | 145 (67%) | 122 (65%) |
| Macroscopic invasion | 158 (37%) | 72 (33%) | 65 (35%) |
| Macrovascular/Extrahepatic | 320 (74%) | 159 (73%) | 136 (72%) |
| BCLC stage | | | |
| B | 140 (32%) | 72 (33%) | 65 (35%) |
| C | 292 (68%) | 145 (67%) | 123 (65%) |
| Alpha-fetoprotein | | | |
| <400 ng/mL | 286 (66%) | 135 (62%) | 123 (65%) |
| ≥400 ng/mL | 146 (34%) | 82 (38%) | 65 (35%) |
| ALBI grade | | | |
| 1 | 142 (33%) | 81 (37%) | 64 (34%) |
| 2 | 290 (67%) | 136 (63%) | 124 (66%) |

Data are presented as median [IQR] or n (%).

Abbreviations: ALBI, albumin–bilirubin index; BCLC, Barcelona Clinic Liver Cancer; ECOG, Eastern Cooperative Oncology Group; HBV, hepatitis B virus; HCV, hepatitis C virus.

Data adapted from *The Lancet Gastroenterology & Hepatology*:

Yau T, Park J-W, Finn RS, et al. *Cabozantinib plus atezolizumab versus sorafenib for advanced hepatocellular carcinoma (COSMIC-312): final results of a randomised phase 3 study.* Lancet Gastroenterol Hepatol. 2023;8(6):502–515. ©2023 Elsevier. Adapted with permission.

### Supplemental Table 3

**Summary of Post-Protocol Therapy and Time to First Systemic Treatment Following Study Drug Discontinuation**

Use of Post-Protocol Therapies

| Category | Cabozantinib + Atezolizumab(N=432) | Single-agent Cabozantinib(N=188) | Sorafenib(N=217) |
|---|---|---|---|
| Systemic therapy use (%) | 111 (26%) | 64 (34%) | 91 (42%) |
| Local therapy use (%) | 7 (1.6%) | 4 (2.1%) | 3 (1.4%) |
| Unknown therapy (%) | 3 (0.7%) | 0 | 0 |

Time to First Post-Protocol Systemic Therapy (months)

| Metric | Cabo + Atezo | Cabo mono | Sorafenib |
|---|---|---|---|
| n (number of patients) | 111 | 64 | 91 |
| Median (range) | 8.31 (0.10–23.42) | 7.41 (0.92–17.08) | 4.43 (0.07–22.96) |
| 25th–75th percentile | 5.85–12.55 | 4.37–10.07 | 2.10–7.79 |

The median time to first post-protocol systemic therapy was longest in the Cabozantinib plus Atezolizumab group (8.31 months), compared to single-agent Cabozantinib and Sorafenib.

### Supplemental Table 4

**Baseline demographics and liver enzyme profiles of patients treated with Cabozantinib or Sorafenib after propensity score matching.**

| Variables | | Overall<br>N = 722 | Cabozantinib<br>N = 361 | Sorafenib<br>N = 361 | p-value |
|---|---|---|---|---|---|
| Age (median [IQR]) | | 65.00 [57.00, 73.00] | 63.00 [56.00, 72.00] | 67.00 [59.00, 74.00] | 0.003 |
| Ethnic (%) | | | | | 0.471 |
| | Non_Hispanic | 669 (92.7) | 339 (93.9) | 330 (91.4) | |
| | Hispanic | 8 ( 1.1) | 3 ( 0.8) | 5 ( 1.4) | |
| | Unknown | 45 ( 6.2) | 19 ( 5.3) | 26 ( 7.2) | |
| Gender (median [IQR]) | | | | | 0.355 |
| | Female | 266 | 139(38.5) | 127(35.2) | |
| | Male | 456 | 222(61.5) | 234(64.8) | |
| Race (%) | | | | | 0.013 |
| | Asian | 29 ( 4.0) | 8 ( 2.2) | 21 ( 5.8) | |
| | Black | 33 ( 4.6) | 14 ( 3.9) | 19 ( 5.3) | |
| | Other | 16 ( 2.2) | 7 ( 1.9) | 9 ( 2.5) | |
| | Unknown | 19 ( 2.6) | 5 ( 1.4) | 14 ( 3.9) | |
| | White | 625 (86.6) | 327 (90.6) | 298 (82.5) | |
| ASTgrade (%) | | | | | **0.01** |
| | 0 | 316 (43.8) | 140 (38.8) | 176 (48.8) | |
| | 1 | 322 (44.6) | 183 (50.7) | 139 (38.5) | |
| | 2 | 53 ( 7.3) | 21 ( 5.8) | 32 ( 8.9) | |
| | 3 | 26 ( 3.6) | 14 ( 3.9) | 12 ( 3.3) | |
| | 4 | 5 ( 0.7) | 3 ( 0.8) | 2 ( 0.6) | |
| ALTgrade (%) | | | | | **0.001** |
| | 0 | 421 (58.3) | 184 (51.0) | 237 (65.7) | |
| | 1 | 235 (32.5) | 137 (38.0) | 98 (27.1) | |
| | 2 | 44 ( 6.1) | 26 ( 7.2) | 18 ( 5.0) | |
| | 3 | 17 ( 2.4) | 10 ( 2.8) | 7 ( 1.9) | |
| | 4 | 5 ( 0.7) | 4 ( 1.1) | 1 ( 0.3) | |
| ASTelevation (%) | | 406 (56.2) | 221 (61.2) | 185 (51.2) | **0.009** |
| ALTelevation (%) | | 301 (41.7) | 177 (49.0) | 124 (34.3) | **<0.001** |

Abbreviations:

AST, aspartate aminotransferase; ALT, alanine aminotransferase

### Supplemental Table 5

**Association between baseline clinical factors and the likelihood of Cabozantinib-induced liver injury**

| Variables | OR (95% CI) | *p*-value |
|---|---|---|
| Race | 1.048 (0.808-1.359) | 0.724 |
| Age | 0.999 (0.978-1.019) | 0.900 |
| Gender | 0.810 (0.518-1.264) | 0.353 |
| ALBI | 0.175 (0.021-1.448) | 0.106 |
| Ethnic | 0.620 (0.375-1.025) | 0.062 |
| APRI | 3.091 (1.011-9.451) | 0.048 |

Abbreviations:
OR, odds ratio; CI, confidence interval; APRI, AST-to-platelet ratio index; ALBI, albumin–bilirubin index; AST, aspartate aminotransferase.

### Supplemental Table 6

**Summary of cabozantinib-induced hepatotoxicity reported in clinical trials**

| Trial | Year | Type of Cancer | Cabo or Cabo + ICBs | | Comparator | |
|---|---|---|---|---|---|---|
| | | | AST increase | ALT increase | AST increase | ALT increase |
| COSMIC-312 | 2021 | HCC | 30.0% | 30.0% | 14.0% | 10.0% |
| COSMIC-021 | 2021 | RCC | 25.3% | 28.1% | 10.9% | 8.4% |
| CELESTIAL | 2018 | HCC | 22.0% | 17.0% | 11.0% | 5.0% |
| NCT00940225 | 2017 | HCC | 27.0% | N/A | 10.0% | N/A |

Abbreviations:

AST, aspartate aminotransferase; ALT, alanine aminotransferase; ICB, immune checkpoint blockade; HCC, hepatocellular carcinoma; RCC, renal cell carcinoma; N/A, not available.
